# Centrosome maintains the integrity of and repairs mature olfactory cilia in adult *Drosophila*

**DOI:** 10.64898/2026.09.02.748762

**Authors:** Minita Desai, Pranjali Priya, Shriya, Haneef Ahmad Dar, Pratik Choudhuri, Pilar Okenve-Ramos, Amit Morarka, Nirpendra Singh, Mónica Bettencourt-Dias, Swadhin Chandra Jana

## Abstract

Mechanisms of cilium assembly are well studied; however, how mature metazoan cilia maintain their structures and functions *in vivo* remains unknown. It is also unclear whether the centrosome-derived basal body (BB) directly contributes to ciliary homeostasis.

We combined biochemistry, genetics, high-resolution subcellular imaging, and electrophysiology to investigate long-lived ciliated olfactory sensory neurons (OSNs) and their olfactory behaviour in adult *Drosophila*. Several centrosome assembly proteins are absent, but another subset persists through dynamic protein exchange at the BBs of mature olfactory cilia. At this ciliary base, pericentriolar material (PCM) components, e.g., γ-Tubulin23C, centrosomin, and pericentrin-like protein form an interconnected network required for ciliary maintenance. Adult- and OSN-specific depletions of these proteins, particularly in combination, disrupt accumulation of the heterotrimeric kinesin-2 at the ciliary base and, subsequently, in the shaft, causing loss of ciliary tubulin, EB1, and odorant receptor co-receptor. These dysregulations cause profound loss of ciliary shaft and impair olfactory function/behaviour. The centrosomal kinases PLK1/POLO and Aurora A also work with this PCM network and are required for mature ciliary homeostasis. Remarkably, these defects in ciliary structure, function, and adult olfactory behaviour are reversible, suggesting that this centrosome-derived BB is a dynamic homeostatic epicentre that regulates trafficking and ciliary structural and compositional integrity *in vivo*.

These findings uncover an active, centrosome-dependent, cell-autonomous mechanism for maintaining and repairing mature metazoan cilia in fully grown organs in adulthood and also provide a possible explanation for how deregulation of conserved ciliary base components could lead to progressive, late-onset cilia-related disorders.

## Introduction

The centrosome, a major cytoskeletal organising centre of eukaryotic interphase cells, forms when centrioles duplicate and acquire a protein-rich matrix called the pericentriolar material (PCM). In a cycling cell, centrosomes play a vital role in spindle assembly before and during cell division. Upon cell cycle exit (G_0_), in some cells they convert into cilia, cells’ signalling hubs with receptors on their membranes (reviewed in^1^). Every cilium has a “base” (centrosome-derived Basal Bodies (BBs) and a Transition Zone (TZ)) and a “shaft” (axoneme, ciliary MicroTubules (MTs), and the signalling receptors on the surrounding membrane)^1,2^. The unique biochemical composition of the ciliary shaft differs from that of the cytoplasm. Therefore, their assemblies mostly rely on various active transports, like kinesin-2/dynein-mediated IntraFlagellar Transport (IFT)-dependent and kinesin-2/dynein-independent traffic (recently reviewed in^3,4^). Cilia mediate external stimulus reception in many unicellular animals and plants, and in all known metazoans. These can be motile (involved in cell motility or causing fluid flow around the cell) or non-motile^5,6^.

Cilia’s vital importance comes into sharp focus through ciliopathies, a diverse group of disorders rooted in deregulation of cilia assembly, structure, or function (reviewed in^5,7,8^). While defects in cilia assembly are known to trigger various diseases, they cannot account for all symptoms observed in conditions like neurodegeneration, retinitis pigmentosa, nephronophthisis, and Alström syndrome, which are marked by gradual cell/tissue decline or late-onset symptoms that emerge later in life^9^. Maintenance of the integrity of all organelles and non-membrane-bound compartments is essential for long-lived cells, which live from a few days to several decades in metazoans. Some organelles, like mitochondria, endoplasmic reticulum, lysosomes, and golgi are considered dynamic^5^. In contrast, cilia have been viewed as static after formation, based on observations like colchicine’s failure to disrupt axonemal MTs, suggesting little or no turnover^10^. But, many ciliated cells, including photoreceptors, sensory neurons, and epithelial cells, persist for years; this raises contradicting and an intriguing possibility that failures in ciliary maintenance drive these diseases. Moreover, work on ciliated cells, including those in *Chlamydomonas* and *C. elegans* first suggested that tubulin is incorporated at the ciliary tip^11,12^. Later, studies in *Trypanosoma*, *Chlamydomonas*, *C. elegans*, and other organisms sporadically explored mechanisms of ciliary shaft homeostasis^13–16^. Therefore, current findings hint that the preservation of metazoan ciliary structure and sensory roles is orchestrated by elusive, cell- and organism-specific mechanisms that remain largely unknown^9^.

Centrosomes have also long been viewed as stable structures due to their resistance to cold and MT-destabilising agents and their stable inheritance across cell cycles^17^. However, during oogenesis in many metazoans, centrosomes are lost from female oocytes and reintroduced in the egg by the male gamete after fertilisation^18,19^. While *C. elegans* centrioles are lost from the ciliary base shortly after ciliogenesis begins^20,21^, the ciliary base is perceived to vary less as *Drosophila*, mice, and human centrioles persist life-time in long-lived ciliated cells. These facts suggest no single rule on centrioles’ or PCM’s stability fits all. Intriguingly, the PCM is assumed to be present in all ciliated cells and mutations in evolutionarily conserved genes encoding ciliary base components display ciliopathies with late-onset symptoms, like Alström syndrome, late-onset brain and endocrine symptoms, and in some patients of retinal degeneration, Joubert syndrome and others^5,7,8^. Nonetheless, centrosomes are probably not intrinsically stable as was thought even a decade ago (reviewed in^22^). A comprehensive description of BB composition and dynamics throughout the lifespan of long-lived ciliated cells is lacking. Moreover, due to lack of studies on centrosome protein deregulation in fully formed cilia, centrosome’s direct role in ciliary homeostasis in any metazoan remains unknown.

In *Drosophila melanogaster*, a genetically tractable model organism, adults harbour Type-I ciliated Olfactory Sensory Neurons (OSNs) in the third antennal segment on their heads. Replenishment of these neurons is minimal (∼1.6% until 15 days of adult life), indicating that ∼1200 ciliated neurons in this tissue persist for the animals lifespan^23,24^. ∼420 sensory hairs, termed sensilla, are present on the surface of the third antennal segment (Figure 1A(ii)). The olfactory cilia innervating these sensilla detect various external odours, which can be quantified through electrophysiological responses (ElectroAntennoGram, EAG) to specific stimuli and behavioural assays^25–28^. Based on morphology, sensilla are broadly classified into three types: basiconic, coeloconic, and trichoid. Basiconic sensilla are further subdivided by size and shape into large (LB), thin (TB), and small (SB) categories^29^. Each OSN extends a single cilium from the distal tip of the dendritic knob. Typically, 2-4 OSNs project their entangled, grouped, ∼15 μm-long cilia into LB sensilla located at the base and medial face of the third antennal segment^27–29^ (depicted in Figure 1A(iii)). External odours pass through the pores present on the sensilla wall and are recognised by the Odorant Receptors (OR) on the ciliary membrane; a major fraction (∼70%) of these different classes of ORs form heterodimeric complexes with Odorant Receptor CO-receptor (ORCO)^30^. Therefore, ORCO’s localisation on the ciliary shaft is a molecular indicator of functional olfactory cilia in *Drosophila*. The olfactory cilia that innervate the large basiconica are the most elaborate and are involved in sensing food-related odours. These cilia grow over 90 hours during pupal development and become fully functional by 2 days of adulthood^27,28^ (depicted in Figure 1A(i)).

**Figure 1:**
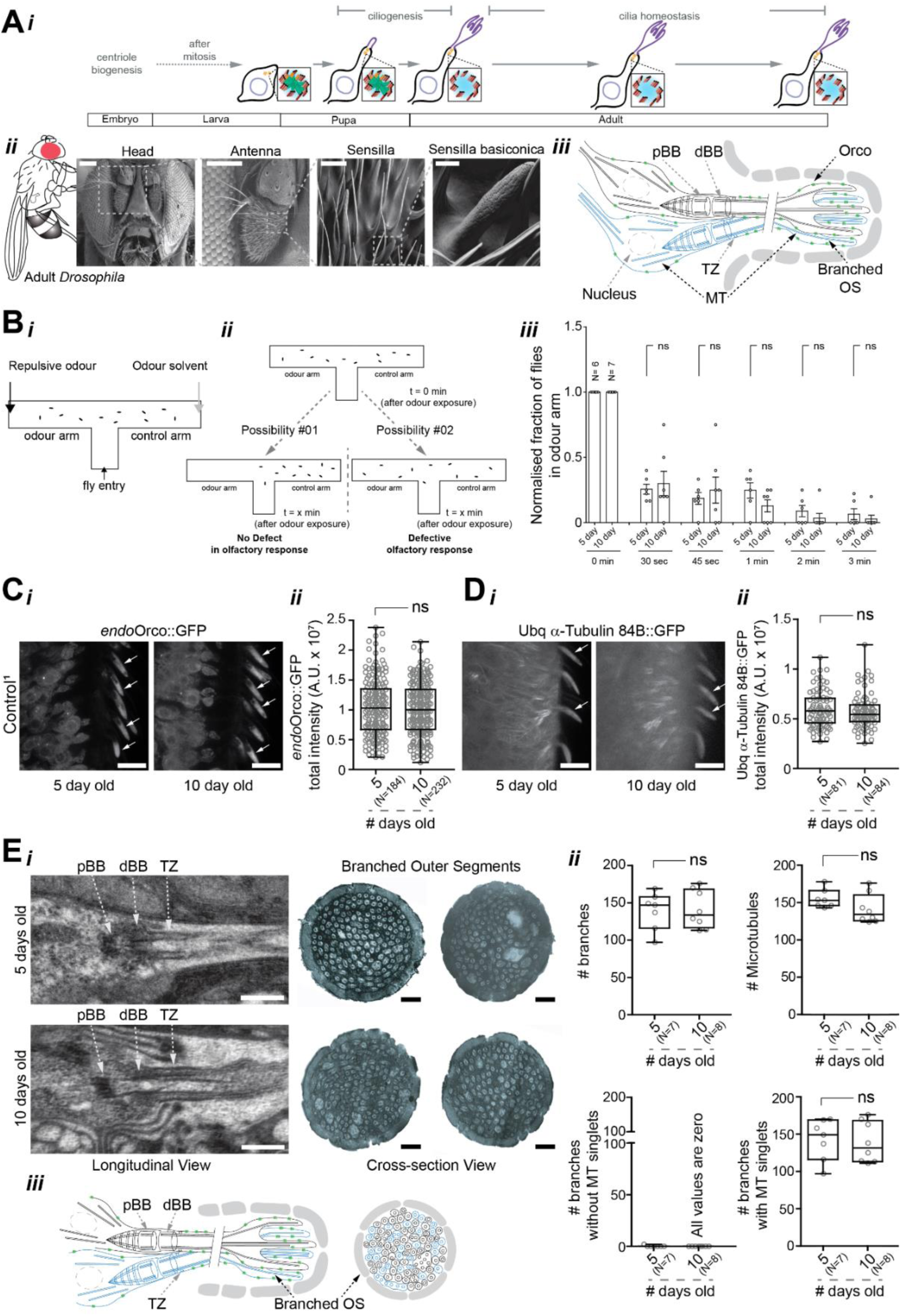
*Drosophila* adult basiconic olfactory cilia maintain their structure and function during adult stage and they are an excellent model to study cilia maintenance in metazoans. A) Ciliogenesis in *Drosophila* OSNs starts during pupal development and completes before fly eclosion^27,28^, making adult flies an excellent model to study cilia homeostasis (i). SEM micrographs of the *Drosophila* head, whole antenna, sensilla on third antennal surface, and individual basiconic sensillum with porous wall (ii). Olfactory cilia are extensively branched in the OS^27,28^ and 2-4 such cilia can innervate a large basiconic sensillum (iii). Odorant receptors on the ciliary membrane form a heterodimeric complex with ORCO for odour sensing^30^. B) Scheme of T-maze odour repulsion assay to assess the olfactory response of fly to a standardised concentration of repulsive odour (i). Flies with no olfactory defect move away from odour arm upon repulsive odour exposure (possibility #1) while flies with defective olfactory response fail to sense the repulsive odour (possibility #2) (ii). Time-dependent changes in the odour repulsion behaviours of 5- day and 10-day-old control adult flies. Each bar corresponds to total of ≥60 flies measured in sets of 7- 10 animals each (iii). C, D) Time-dependent changes in intensity of *endo*ORCO::GFP (C) and *Ubq* α-Tubulin84B::GFP (D) in the basiconic olfactory cilia of 5-day and 10-day-old control adult flies. Representative images (i) and respective quantifications (ii). E) Representative electron micrographs of longitudinal sections (i, left) marking the pBB and dBB, TZ, and cross sections of the branched OS (i, right) from 5-day (top) and 10-day-old (below) control adult flies and respective branch quantifications (ii). Scheme of OSNs innervating basiconic sensilla showing the pBB, dBB, TZ, and branched OS (iii). Scale bars in SEM micrographs in A (ii) are - 100 µm (head and antenna), 5 µm (sensilla) and 2 µm (sensilla basiconica), those in the confocal micrographs in C and D are 10 µm, and those on all TEM micrographs in E are 500 nm. All electron micrographs in E represent features observed in n=3 samples (the experiments were repeated independently at least twice with similar results).

Therefore, here we used a combination of techniques, including biochemistry, genetics, high-resolution subcellular microscopy, electrophysiology, and animal behaviour, to investigate the long-lived ciliated large basiconic OSNs in *Drosophila*. Specifically, we aimed to elucidate three points. First, whether and how much of the BB composition remains consistent and dynamic after these cilia are fully assembled and functionalised (Figure 1-3). Second, the orchestrated role of key ciliary base structural proteins and centrosomal regulatory kinases in maintaining structure and function of the ciliary base and shaft (Figure 4-7). Third, if deregulating ciliary base protein composition disrupts the ciliary shaft structure and function, can the resulting defects be reversed (Figure 8)?

**Figure 2:**
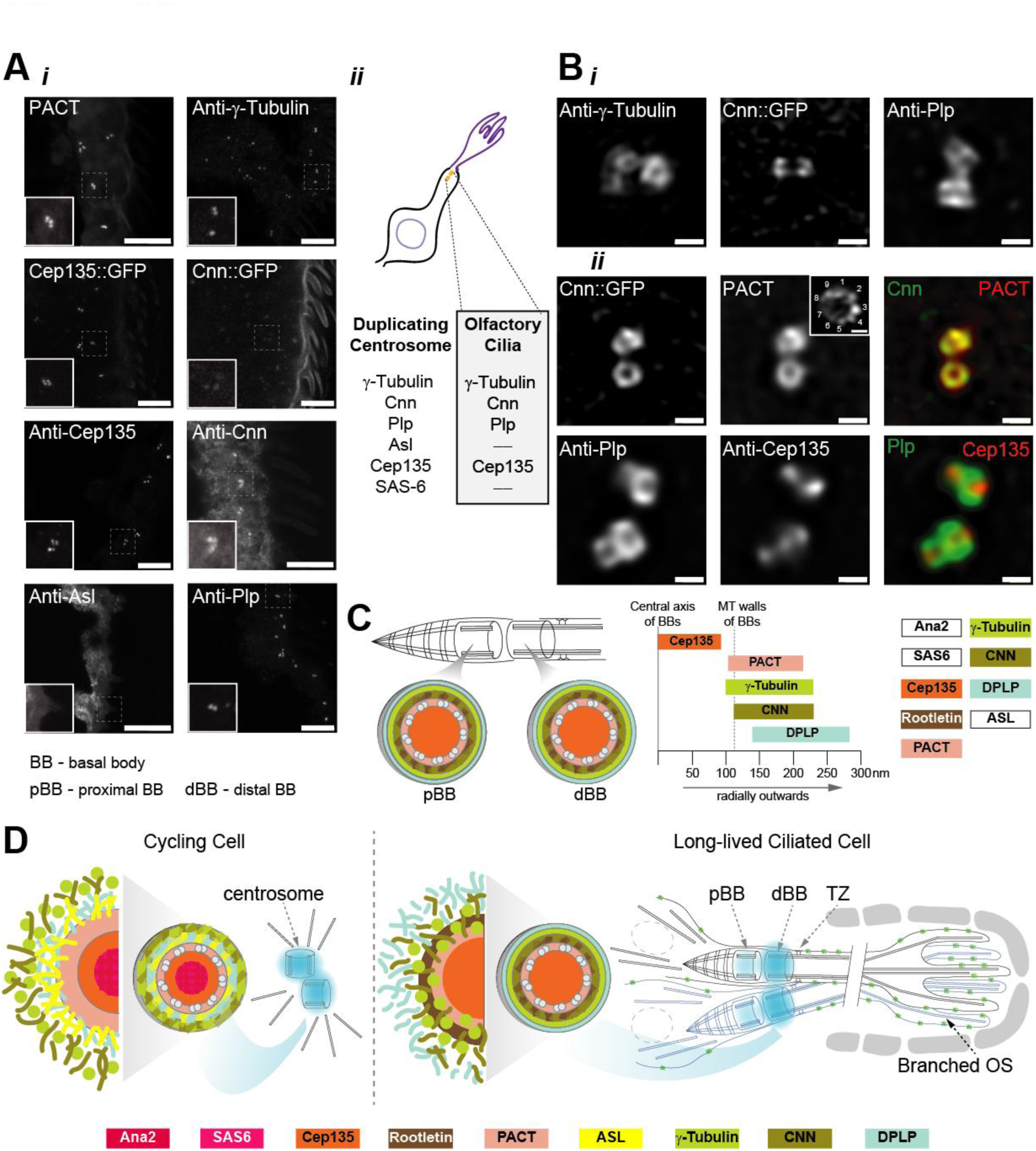
Localisation, organisation and quantification of localisation of endogenous centrosome proteins in basiconic olfactory cilia of 5-day old adult antenna. A) Representative confocal images of the four centrosomal proteins (γ-Tubulin, CNN, PLP, CEP135) and centriole wall marker (PACT) present in adult *Drosophila* antennal tissue, while ASL, a key PCM expansion factor in cycling cells, is absent (i). List of centriole and PCM proteins present in duplicating centrosome and olfactory cilia of *Drosophila* (ii). B) Representative Airyscan 2.0 images describe the localisation of four centrosomal proteins (γ-Tubulin, CNN, PLP, CEP135) and centriole wall marker PACT used as reference in olfactory cilia (i, ii). UxM micrograph of cross section showing 9-fold localisation of PACT::GFP at the cilia base (inset in ii). C) The schemes representing the localisation patterns of the proteins in the pBB and dBB cross sections (left) and relative localisation spread of each centrosome protein (right). Quantifications of localisation is described in Supplemental Figure 02. PCM protein ASL (shown here) and cartwheel proteins ANA2 and SAS6 (shown previously^2^) were not detected in olfactory cilia. CNN localises closer to BB walls while γ-Tubulin and PLP form radially outward and thicker diameter rings. D) Scheme comparing the centrosome of cycling cells and BBs of long-lived ciliated cell. PCM protein composition and their organisation at the OSN ciliary base is different from the cycling cells. The experiments presented in A and B were repeated independently thrice. Scale bars on micrographs in A and B are 10 µm and 500 nm, respectively, while that in B (ii, PACT inset) is 100 nm.

**Figure 3:**
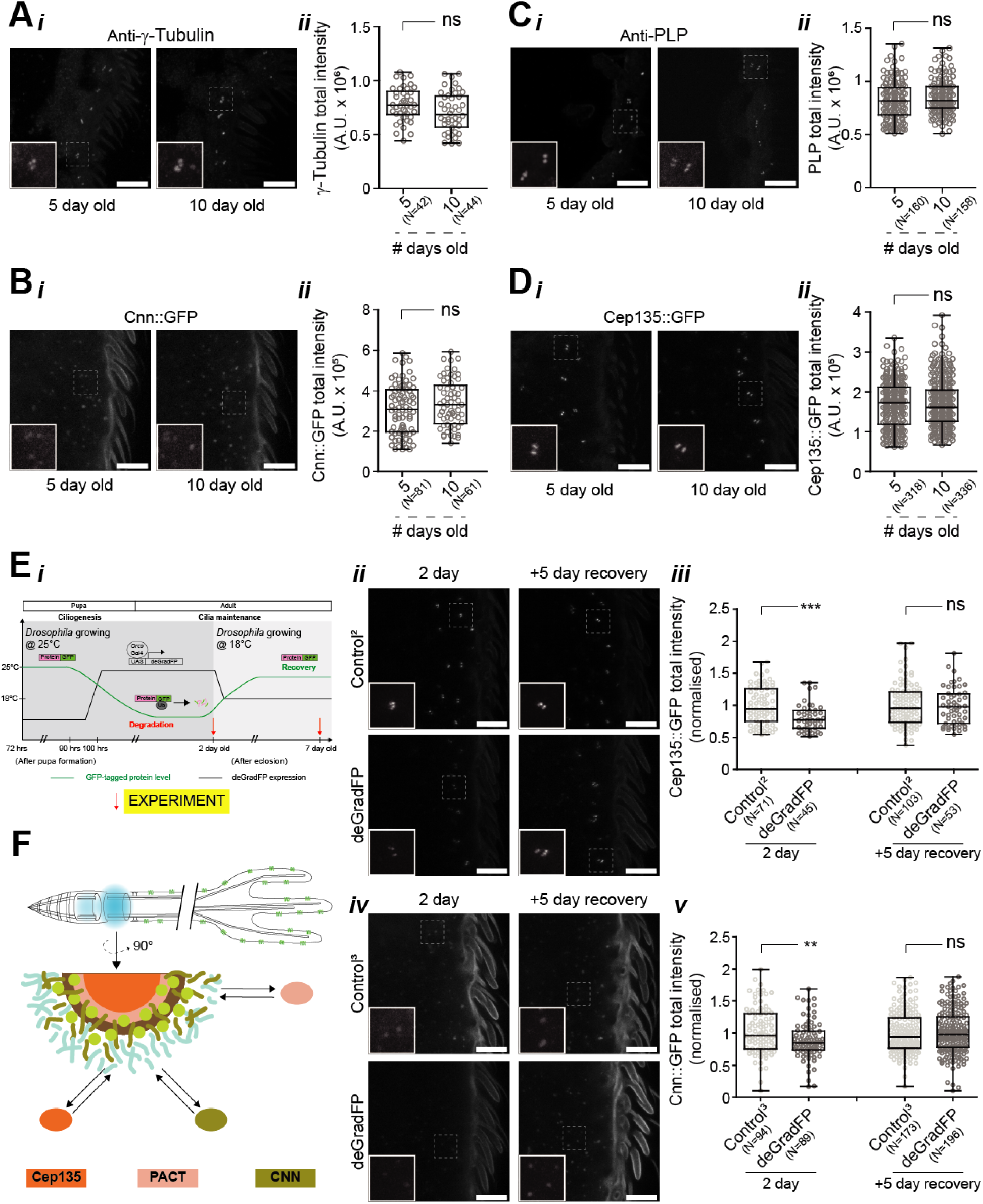
Centriole and PCM proteins remain present till at least 10 days and continue to exchange dynamically at the base of basiconic olfactory cilia during the adult stage. A-D) Time-dependent changes in intensity of four centrosomal proteins - γ-Tubulin (A), *endo*CNN::GFP (B), PLP (C), and *endo*CEP135::GFP (D) in basiconic olfactory cilia of 5-day and 10-day-old control adult flies. Representative images (i) and their respective quantifications (ii). The selected centrosomal proteins that persist at the ciliary base in adult olfactory cilia are maintained till 10 days of age. E) Scheme of the approach and timeline of the conditional degradation experiments (i). Time-dependent changes in intensity of centriolar protein *endo*CEP135::GFP (ii, iii) and PCM protein *endo*CNN::GFP (iv, v) upon conditional degradation (at 25°C under Gal4*^Orco^* driver) for 2 days and recovery (at 18°C) for 5 days post-degradation. Representative images (ii, iv) and their respective quantifications (iii, v). F) Scheme depicting the molecular organisation of centrosomal proteins at the ciliary base of long-lived ciliated OSN and showing PACT (see Supplemental Figure 03), CEP135 and CNN (shown in E) are dynamically exchanged at the base during cilia maintenance of adult flies. Scale bars on all the confocal micrographs are 10 µm. All-range box plots of total intensity of GFP- tagged proteins at the base of olfactory basiconic cilia in the flies with specific genotypes and specific experimental condition are stated on the graphs.

**Figure 4:**
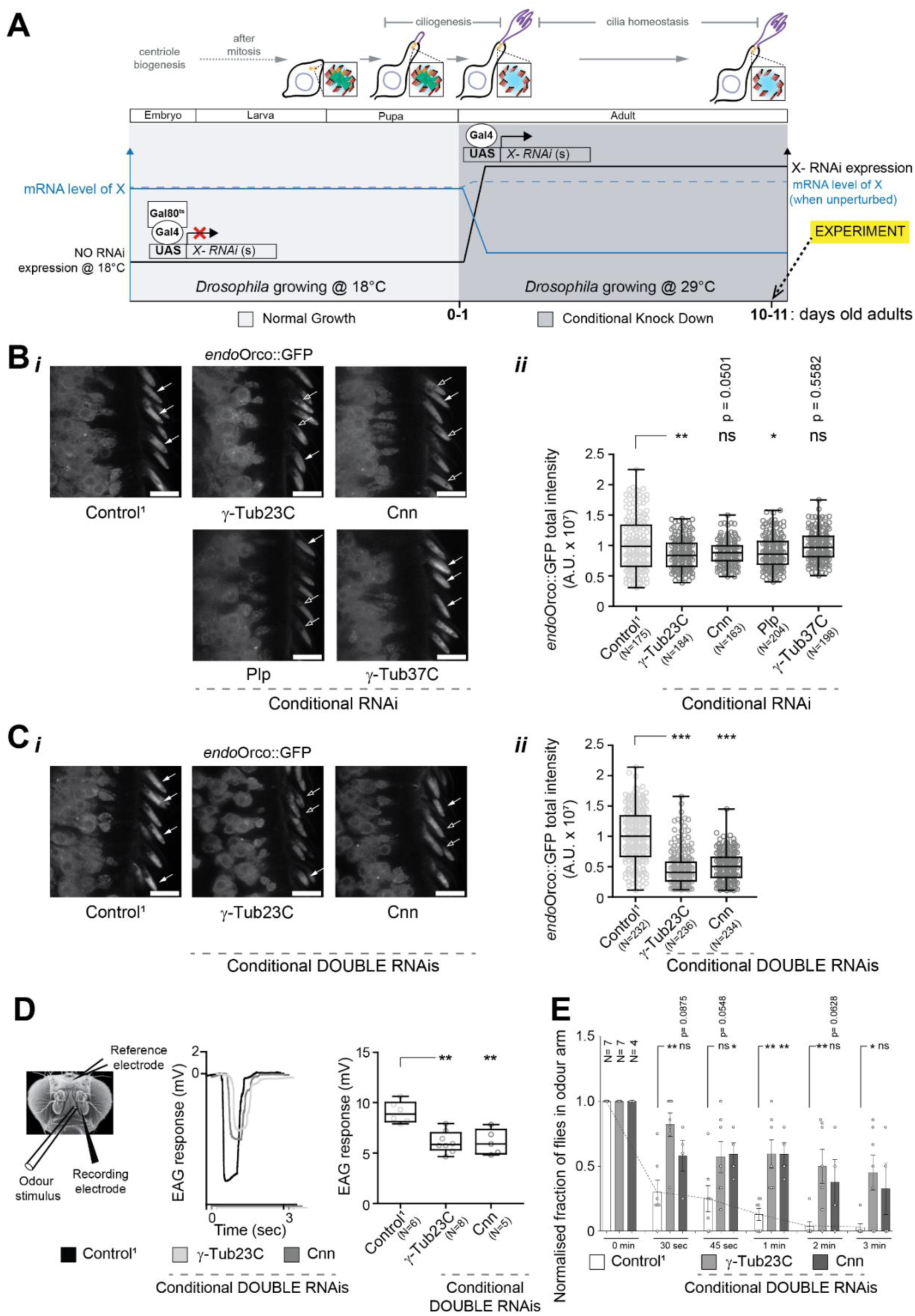
γ-Tubulin23C, Centrosomin (CNN), and Pericentrin-like protein (PLP) are essential for the maintenance and structure of basiconic olfactory cilia, whereas γ- Tubulin37C is not required. A) Scheme of the approach and timeline of the conditional knockdown experiments using OSN-specific Gal4*^Orco^* temporally regulated by temperature-sensitive Gal80^ts^. RNAi expression was suppressed during development to allow for ciliogenesis in OSNs. Upon eclosion, the flies were shifted to 29°C to induce knockdown of selected centrosomal proteins in the adult OSNs. Effect of the knockdown on ciliary structure and function was studied on 10-day-old adult flies. B) Changes in *endo*ORCO::GFP intensity in the basiconic olfactory cilia of 10-day-old control flies and flies with conditional knockdown of γ-Tubulin23C, γ-Tubulin37C, CNN, and PLP. Representative images (i) and respective quantifications (ii). C) Changes in *endo*ORCO::GFP intensity in the basiconic olfactory cilia of 10-day-old control flies and flies with conditional DOUBLE RNAi knockdown of γ-Tubulin23C and CNN. Representative images (i) and respective quantifications (ii). D) Scheme of electrode position and odour delivery on immobilised fly head for recording electrophysiological response (left). EAG response was measured from basiconic region of third antenna of 10-day-old control flies and flies with conditional DOUBLE RNAi knockdown of γ-Tubulin23C and CNN. EAG traces (middle) and respective quantifications (right). E) Time-dependent changes in the odour repulsion behaviours in 10-day-old control flies and flies with conditional DOUBLE RNAi knockdown of γ-Tubulin23C and CNN. Each bar corresponds to a total of ≥60 flies measured in sets of 7-10 animals each. Scale bars on confocal micrographs in B and C are 10 µm.

## Results

### Olfactory basiconic cilia structure and function remain intact in adult flies at least up to 10-days age

This study requires the selection of a cell type that is both ciliated and long-lived (Figure 1A(i)). The sensilla subtype (LB-I) examined in this study is developmentally defined and numbers ∼15 sensilla per antenna, with consistent location across individual adult antenna^29^. This consistency enables precise spatiotemporal analysis of these cilia and their function. Olfactory reception behaviour in 5-day-old flies is comparable to that observed in adult flies when they achieve full odour reception functionality (Figure 1B(iii)).

To begin with, we assessed the olfactory cilia function using an odour-repulsion assay in a T-maze, where wild-type adult flies were exposed to a repellent odour (e.g., Benzaldehyde) from one arm. Wild-type flies typically move away from the repellent odour, resulting in fewer flies in the repulsive odour arm over time^26^. Our results indicate that 5-day-old and 10-day-old flies display similar odour-repulsion behaviour (Figure 1B). Given that sensory behaviour defects increase with age at 25°C, ciliary structure and the localisation of key proteins were examined in 5- and 10-day-old flies. We found cilia length, ∼15μm, remains stable between these ages (Supplemental Figure 1A). Levels of endogenous ORCO::GFP (referred as *endo*ORCO::GFP or ORCO here) at the ciliary shaft, and α-Tubulin84B, a major tubulin isoform of the ciliary skeleton, also remain unchanged (Figure 1C, D). We then analysed the ultrastructure of these olfactory cilia, previously described in 1-day-old flies^2,27,28^, in 5- and 10-day-old flies. Consistent with previous reports for 1-day-old flies, two BBs are linearly arranged and connected by a rootlet also in 5- day-old flies (Figure 1E(i) (left) and Supplemental Figure 1C). Like duplicating centrioles, MTs within these BBs display a 9-fold symmetry, and the electron-dense material, likely the PCM, is observed between the MTs and along the BB wall (Supplemental Figure 1D). The distal BB at the end of the dendritic knob forms a TZ. All these features are present in both 5- and 10-day-old flies (Figure 1E(i) (left) and Supplemental Figure 1C). Additionally, numerous singlet MTs extend from the distal end of the TZ, and each singlet MT at the distal end of the cilia, known as the Outer Segment (OS), is surrounded by ciliary membranes (Figure 1E(i) (right), (iii), and Supplemental Figure 1B). At the OSs, the total number of branches (5-day: 139±25 and 10-day: 141±25), branches containing MTs (5-day: 142±28 and 10-day: 139±27), and singlet MTs (5-day: 155±13 and 10-day: 141±20) remains consistent between 5- and 10-day-old flies (Figure 1E(ii)). These findings suggest that ciliary shafts and bases, including membrane and skeletal components, are maintained either as stable or as dynamic skeleton structures in both 5- and 10-day-old flies. This system is therefore perfectly suited for investigating whether ciliary bases, in particular BBs, contribute to ciliary homeostasis *in vivo* and elucidating the underlying mechanisms.

### Composition and organisation of PCM proteins at the olfactory ciliary base show changes compared to the duplicating centrosome, and persistent proteins continue to be actively replenished

Consistent with centrosomes, the BBs of olfactory cilia in 5- and 10-day-old flies are surrounded by electron-dense material (Figure 1E(i) (left) and Supplemental Figure 1C, D). To investigate the roles of centrosomal proteins in ciliary homeostasis, we first assessed whether the composition of ciliary base, particularly centriole and PCM proteins, in young adult flies aligns with composition and organisation observed in *Drosophila* cycling cells^31,32^. We examined the localisation of several evolutionarily conserved centriole proteins (SAS6, STIL/SAS-5/ANA2, CEP135/BLD10, PACT) and four PCM proteins (TUBG1/TBG-1/γ-Tubulin, CDK5RAP2/SPD-5/CNN, PCNT/PCMD-1/PLP, CEP152/ASL) at the centrosome in cycling cells (Supplemental Figure 2A) and at the ciliary base of OSNs in 5-day-old adult antennae (Figure 2A(i)). Based on reagent availability, fluorescent-tagged candidate centrosomal proteins expressed under endogenous promoters or antibodies against candidate proteins were used (protocols described in^33^; see “Materials and methods” section for details). All centriole and PCM proteins examined were detected in cycling cells (Supplemental Figure 2A, Supplemental Table 01, and previously described in^31,32^), but several markers were not observed at the ciliary bases of 5-day-old OSNs (Figure 2A(i, ii)). While CEP135/BLD10, γ- Tubulin, CNN, and PLP persist at the BBs, SAS6, ANA2, and ASL are absent from the olfactory cilia BBs in 5-day-old flies (Figure 2A(i, ii)). This result is consistent with previous report using similar markers in the olfactory cilia in 0-1-day-old flies^2^. The absence of ASL at the BBs of olfactory basiconic cilia indicates that this PCM is likely unable to template a new centriole, and that PCM proteins composition here differs from that in cycling cells. However, the presence of key PCM structural proteins, such as γ-Tubulin, CNN, and PLP, suggests that these BBs continue to function as MT-organising centres (MTOCs), as cytoplasmic MTs are observed surrounding them in both flies and other species^2,20,21^. Then the relative localisation order of PCM proteins with respect to CEP135, an inner centriole wall component, was investigated at the ciliary base using super-resolution imaging. PACT (Pericentrin-AKAP450 Centrosomal Targeting) domain localisation corresponds to the centriole wall dimensions estimated from TEM cross-section images (Supplemental Figure 1D) and super-resolution images collected using expanded antennal sections (Figure 2B(ii)). PCM organisation around these BBs differs from that of centrosomes in *Drosophila* S2 cells^31,32^ (depicted in Figure 2D). Super-resolution imaging indicates that the localisation of γ-Tubulin, CNN, and PLP around the BB walls partially overlaps (Figure 2B, C and Supplemental figure 2B, C). Unlike PCM organisation in S2 cells, CNN localises closer to the BB walls, while γ-Tubulin and PLP form radially outward and thicker diameter rings (Figure 2B, C and Supplemental figure 2B, C). CNN, γ-Tubulin, and Pericentrin have also been identified at many ciliary bases in other organisms^20,21,34,35^. These altogether suggest potential combinatorial roles for these three proteins in the PCM regions surrounding the BBs in fully assembled and functional olfactory cilia.

The persistence of the proteins that we investigated in the previous section at the ciliary bases of 5- and 10-day-old flies was next examined. All centriole and PCM proteins previously found to localise at the ciliary base in adult flies continued to localise and maintain their protein levels even in 10-day-old flies (Figure 3A-D). These findings suggest that these proteins are either components of a static BB structure or are dynamically replenished at the ciliary base. To distinguish between these possibilities, an experiment was designed using the Gal4*^Orco^*- *UAS*deGradFP-*Ubq/endo*X::GFP system, in which deGradFP was spatiotemporally expressed in OSNs of flies expressing GFP-tagged versions of endogenously expressed centrosomal proteins (Figure 3E(i)). X represents a candidate gene/protein. We used the Gal4*^Orco^* driver, which expresses at a very late stage of the cilia maturation in most (∼70%) of OSNs at the third antennal segment and continues to express in adult flies^28,30^. If protein levels at the ciliary base could be reduced using this approach, it would indicate that GFP-tagged proteins localising there are degradable. By two days of adult life, application of Gal4*^Orco^*resulted in a significant reduction in the levels of a domain of PLP, namely *Ubq*-GFP::PACT (expressed using the Tubulin promoter and works as a reference to centriole walls^36^), at the olfactory ciliary base (Supplemental Figure 3A, C). Following two days of ectopic degradation, cessation of deGradFP expression in the same cells led to recovery of PACT::GFP levels to those observed in controls (Supplemental Figure 3B, D). This experiment demonstrates the successful development of a tool to study the dynamic properties of tagged centrosome proteins expressed under an ectopic promoter. Furthermore, when CEP135::GFP (Figure 3E(ii, iii)) and CNN::GFP (Figure 3E(iv, v)) were expressed under their respective endogenous promoters, two days of conditional knockdown in adult flies resulted in reduced protein levels at the ciliary bases. Upon cessation of deGradFP expression, the levels of CEP135 (Figure 3E(ii, iii)) and CNN (Figure 3E(iv, v)) proteins became indistinguishable from their respective controls. Collectively, these results indicate that some centrosomal proteins persisting at the ciliary base are dynamically replenished (depicted in Figure 3F).

### PCM structural proteins, γ-Tubulin23C and CNN, work together at the ciliary base to maintain ciliary shaft’s shape and function

All three PCM proteins (γ-Tubulin, CNN, and PLP) that we investigated are also essential for expansion and maturation of duplicating centrosome. Consequently, mutant flies lacking these genes, thus proteins, exhibit defective ciliogenesis. As a result, previously generated mutants could not be utilised for this study on cilia homeostasis. Instead, we developed tools enabling cell (OSN)- and time (adult)-specific control of gene expression (Figure 4A).

We first tested the efficacy of the RNAi lines (with a hairpin targeting an exon common to most isoforms of a candidate gene and without any predicted off-targets) using a ubiquitously active promoter (Gal4*^Tubulin^*; see Supplemental Figure 4A) (knockdown efficiencies observed by us are in the range of 60-99%, Supplemental Figure 4C, 7A(iv)). We found that the mRNA levels of most isoforms are strongly reduced (≥60%) in the flies expressing the hairpins of candidate genes (γ-Tubulin23C, γ-Tubulin37C, CNN, and PLP) using Gal4*^Tubulin^*, as compared to control (Supplemental Figure 4C). Importantly, the larval-to-pupae development is significantly reduced (Supplemental Figure 4B(i)), and most of the pupae failed to convert to adult flies in the case of γ-Tubulin23C and PLP individual knockdowns compared to controls (Supplemental Figure 4B(ii)), consistent with the centrosome/cilium defects reported in their respective genes’ mutants^36,37^. In contrast, for CNN knockdown, while no defect was observed in the larva-to-pupa conversion (Supplemental Figure 4B(i)), pupa-to-adult conversion was marginally affected (but not significantly reduced, Supplemental Figure 4B(ii)), and the adult flies showed marginally defective negative geotaxis (Supplemental Figure 4B(iii)). Indeed, the homozygous CNN mutants grow to adulthood, suggesting that the knockdown flies exhibit a phenotype similar to the CNN loss-of-function mutant^38^. Given that γ-Tubulin37C is critical for cell/nuclear divisions till stage 9 of the embryo, γ-Tubulin37C mutant embryos fail to grow^39^. Therefore, role of γ-Tubulin37C in later stages of larvae/pupae/adults is unclear. Using the available hairpin of γ-Tubulin37C, we successfully reduced (≥55%) the mRNA levels (Supplemental Figure 4C(iv)), and the adult flies expressing the hairpin show reduced negative geotaxis ability (Supplemental Figure 4B(iii)), suggesting its role in centrosome/cilia assembly during the fly development. These analyses suggest that our selected RNAi hairpins are appropriate for proceeding with our conditional knockdown experiments.

Then, to test for defects in olfactory cilia maintenance, we used a temperature-sensitive (ts) system that allows us to control gene expression in a OSN- and adult-specific manner (Gal4*^Orco^*-*UAS*X-RNAi-*Tub*Gal80^ts^) (Figure 4A; see the “Materials and Methods” section)^40^. X represents a candidate gene/protein. The *Tub*Gal80^ts^ system (at 18°C, *Tub*Gal80^ts^ is active, thus there is no RNAi knockdown; at 29°C, *Tub*Gal80^ts^ is inactive, thus knockdown through RNAi is active) allowed us to acutely down-regulate candidate genes’ expression with an RNAi line just during adulthood by moving to and rearing flies at 29°C after flies emerge from the pupae (1-day-old flies, Figure 4A). This experimental setup further allows us to completely disentangle the functions of all four genes in centrosome maturation or cell division from their under-investigated role in adult cilia maintenance. ORCO localisation on the ciliary shaft serves as a molecular indicator of functional olfactory cilia in *Drosophila*^30^. To assess this, we quantitatively analysed *endo*ORCO::GFP localisation in the ciliary shaft of large basiconic OSNs (LB-I) in 10-day-old flies. Marginal reductions in ciliary ORCO levels were observed following knockdown of γ-Tubulin23C (reduction by ∼15%), CNN (reduction by ∼13%), and PLP (reduction by ∼13%) individually, whereas no reduction was detected in γ-Tubulin37C knockdown compared to control flies (Figure 4B). These findings suggest that γ-Tubulin37C is unlikely to contribute to olfactory cilia maintenance, while γ-Tubulin23C, CNN, and PLP may each play roles in maintaining ciliary structure and function.

The marginal reduction (∼13-15%) in ciliary ORCO in single-RNAi flies are less assuring of these proteins’ role in cilia homeostasis, thus we explored two hypotheses. First, the use of two distinct hairpins targeting a candidate gene may yield a stronger spatiotemporal knockdown. Second, because the PCM surrounding the centriole is a highly dense protein matrix, using two distinct hairpins targeting two different candidates that interact at the BB may also result in a stronger defect. Assumingly, ciliary ORCO was dramatically reduced in OSNs in γ-Tubulin23C double RNAi (reduction by ∼54%) and CNN double RNAi (reduction by ∼50%) compared to control (Figure 4C). We achieved ∼65% reduction in the γ-Tubulin23C mRNA levels in the antennal tissue of γ-Tubulin23C double RNAi flies (Supplemental Figure 4D). Olfactory cilia function was further evaluated by measuring EAG from the third antennal segment in response to an odour (e.g., ethyl acetate)^25,27^ and by assessing odour repulsion behaviour to a repellent odour (e.g., Benzaldehyde)^26^. Average EAG responses in adult flies with γ-Tubulin23C double RNAi (6.1±1.1 mV) and CNN double RNAi (6.1±1.3 mV) were significantly reduced compared to control (9.1±1.1 mV) (Figure 4D). The observed defects in ciliary ORCO levels and reduced EAG responses indicate that this adult-specific γ-Tubulin23C double RNAi and CNN double RNAi OSNs exhibit defective ciliary membrane composition, resulting in impaired ciliary function. These defects in ciliary functions then lead to impaired odour repulsion behaviour in the double RNAi flies (Figure 4E). Collectively, these results demonstrate the essentiality of γ-Tubulin23C and CNN, independently, in maintaining olfactory reception function after ciliogenesis is completed, supporting the first hypothesis for structural proteins at ciliary bases.

To test the second hypothesis, we aimed to determine whether these proteins interact at the ciliary bases in adult flies, by establishing a technique to purify centrosomes (using modified version of previously described protocol^41^) from embryos, which contain duplicating centrosomes (as a control for comparison), and from the head/antenna, which contain non-duplicating centrosomes or ciliary bases (Figure 5A(i) and Supplemental Figure 5C).

**Figure 5:**
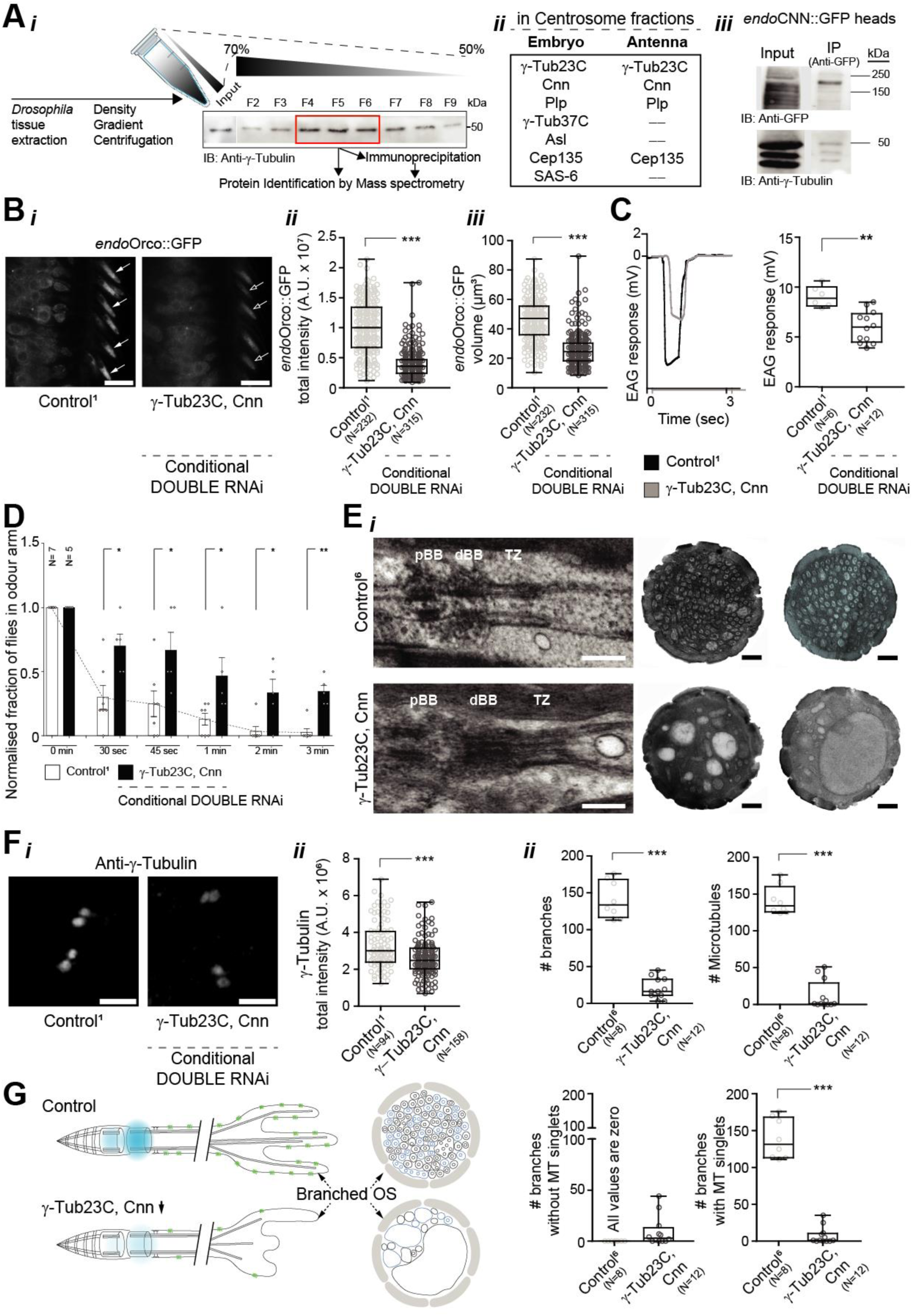
γ-Tubulin23C and CNN physically and genetically interact for maintenance of the structure and function of adult basiconic olfactory cilia. A) Scheme of the approach of protein identification in purified centrosomes from *Drosophila* cycling cells (embryo) and ciliated tissues (adult heads/antenna) (i). List of centriole and PCM proteins found in isolated centrosomal fraction from 2-4-hour-old embryos and whole antenna from 0-2-day-old *endo*CNN::GFP flies (ii). CNN (probed by anti-GFP) and γ-Tubulin both are detected in 0-2-day-old adult fly heads total protein lysate (Input, iii). γ-Tubulin is part of the co-IP complex of *endo*CNN::GFP isolated from heads (IP, iii). B) Changes in *endo*Orco::GFP intensity (Representative images (i) and respective quantifications (ii)) and volume (iii) in the basiconic olfactory cilia of 10-day-old control flies and flies with conditional simultaneous knockdown of γ-Tubulin23C with CNN. C) EAG response measured from basiconic region of third antenna of control and conditional knockdown flies. EAG traces (left) and respective quantifications (right). D) Time-dependent changes in the odour repulsion behaviours of control and conditional knockdown flies. Each bar corresponds to a total of ≥60 flies measured in sets of 7-10 animals each. E) Representative electron micrographs of longitudinal section showing the pBB, dBB, and TZ of control and conditional knockdown flies (i, left). Representative electron micrographs of individual cross sections of ciliary OS (i, right) and branch quantification (ii). F) Changes in intensity of γ-Tubulin in basiconic olfactory cilia in control and conditional knockdown flies. Representative images (i) and respective quantifications (ii). G) Scheme depicting olfactory cilia with extensive OS branching in presence of functional interacting PCM (shown in blue). Removal of PCM components such as γ-Tubulin23C and CNN leads to severe branching defects and MT loss from OS leading to loss of structural integrity of the olfactory cilia. Scale bars on confocal micrographs in B and F are 10 µm and 2.5 µm respectively, while those on all TEM micrographs in E are 500 nm each.

Interestingly, several proteins involved in centrosome duplication, like γ-Tubulin37C, ASL, and SAS6, were absent from the purified ciliary base. In contrast, proteins like γ-Tubulin23C, CNN, PLP, and CEP135 were found in both duplicating centrosomes and ciliary bases fraction (Figure 5A(ii) and Supplemental Figure 5D(ii)), consistent with our protein localisation results (Figure 2 and Supplemental Figure 2). Additionally, γ-Tubulin23C was found to co-immunoprecipitate (IP) with CNN in ciliary base fractions (Figure 5A(iii)), as well as in centrosomal fractions from embryos^42^ (Supplemental Figure 5D(i)). These findings suggest that these proteins highly likely function together as a network at the ciliary base. To further investigate this possibility, flies were generated with OSN- and adult-specific knockdown of two different PCM proteins by expressing combinations of hairpin RNAs targeting two PCM candidates simultaneously. The effects on ciliary defect phenotypes were then assessed. Notably, simultaneous removal of γ-Tubulin23C with CNN resulted in a dramatic reduction of ciliary ORCO (∼62%) and its volume (∼45%) compared to control (Figure 5B). These combination double RNAi flies also exhibited significant defects in EAG responses and odour repulsion responses (Figure 5C, D). Interestingly, combining single RNAis of two different PCM proteins caused more severe ciliary defects (ciliary ORCO reduction by 62%) than double RNAis of a single protein (50-54% reduction). This supports the conclusion that the PCM is stabilised by various proteins, and at least these two (γ-Tubulin23C and CNN) act synergistically at the ciliary base.

Though, at the ciliary bases, the electron dense walls of a few BBs were marginally distorted in all conditional double RNAi flies (∼8% of ciliary bases examined), we found that two BBs are always arranged linearly and have a TZ per ciliary base in the combination double RNAi flies (100%) as observed in the control (Figure 5E(i) (left) and Supplemental Figure 6B). These suggests that BBs at the ciliary base continue to persist when γ-Tubulin23C and CNN are adult-specifically removed from OSNs. But ultrastructural analysis of the ciliary OS using cross-sectional images of basiconic sensilla (LB-I) demonstrates that the characteristic, highly branched finger-like projections, which comprise nearly 141±25 branches with singlet MTs per sensillum in control sensilla, are significantly reduced in double RNAi sensilla. The double RNAi for CNN exhibit a total of 45±30 branches, whereas double γ-Tubulin23C RNAi results in 30±10 branches (Supplemental Figure 6A). When two single RNAis targeting γ-Tubulin23C and CNN are combined, ciliary finger-like projections were nearly abolished (a total of 20±15 branches with or without MTs per sensillum) (Figure 5E(i) (right) and (ii)) and Supplemental Figure 6B). In the combination double RNAi flies, the total number of branches containing singlet MTs is 7±10, suggesting a profound loss of ciliary shaft (a reduction of about 95% compared to control flies; Figure 5E(i) (right) and (ii)) and Supplemental Figure 6B; schematic presentation of defects is depicted in Figure 5G and Supplemental Figure 6C). Also, γ-Tubulin localisation at the ciliary base is reduced (by ∼20%) in the combination double RNAi flies, compared to control flies (Figure 5F). In summary, these results suggest that OSN- and adult-specific PCM removal does not remove centrioles at the ciliary base, but it induces yet unknown changes in ciliary base properties, including the dynamicity of the environment around the BBs. These properties are necessary to maintain ciliary skeleton and membrane structure integrity and ORCO localisation, thereby the ciliary function (see scheme in Figure 5G).

### Molecular mechanism of ciliary shaft maintenance by γ-Tubulin23C-CNN-PLP at ciliary base

We sought to determine how this process is regulated and identify the mechanisms that maintain ciliary base, thus ciliary shaft structure and function. To address this, we studied fluorescent-tagged ciliary base and shaft proteins after OSN- and adult-specific removal of γ- Tubulin23C and CNN combined for 2 days. These analyses would help us identify which process goes awry early and might lead to the dramatic defects observed after 10 days of knockdown. We found that removing PCM proteins via simultaneous knockdown of γ- Tubulin23C with CNN reduced tubulin (α-Tubulin84B::GFP) levels in the ciliary shaft (Figure 6A(i)). Similarly, End-Binding protein 1 (EB1), a MT-plus-end-binding and MT-stabilising protein, was significantly decreased in the ciliary shaft compared to controls (Figure 6A(ii)). These findings strongly suggest that defects in the ciliary shaft MT-skeleton structure likely contributes to impaired ciliary ORCO localisation, altered cilia morphology, and thus compromised cilia function. Consistent with this hypothesis, we previously established that the branching and MT singlets at OS are significantly lost upon PCM components removal (Figure 5E(i) (right) and Supplemental Figure 6). In contrast, analysis of CEP290, a key component of the TZ, and CEP135, a core protein of the centriole/BB, revealed no significant defects in these markers in the combination double RNAi compared to respective controls (Figure 6A(iii, iv)). These observations are consistent with our results described in the earlier section that the ultrastructural organisations of the ciliary base are broadly unaffected upon PCM components removal (Figure 5E(i) (left) and Supplemental Figure 6) further confirming that the core structure of ciliary base remains intact despite the dynamicity of the PCM around the BBs.

**Figure 6:**
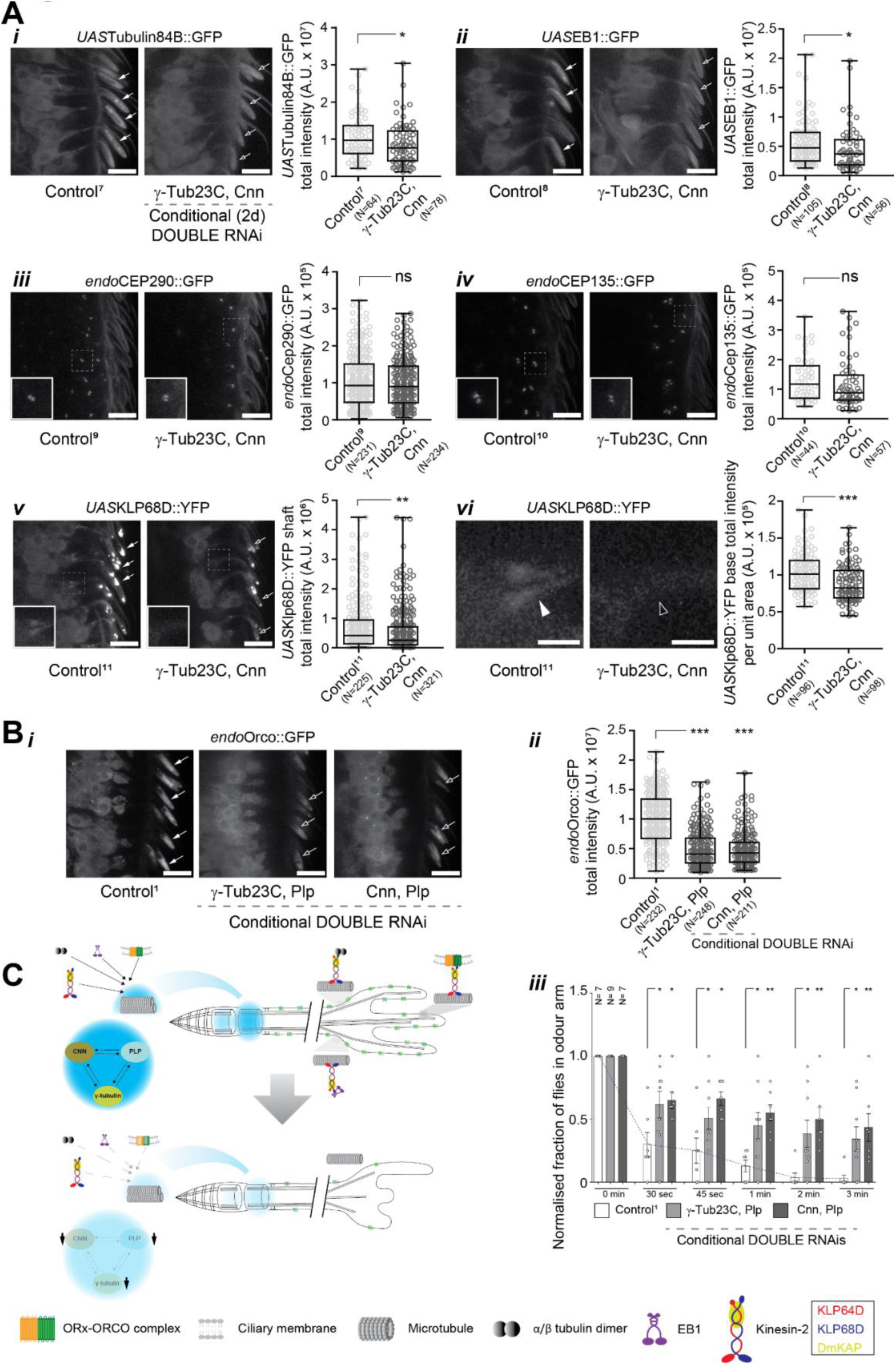
Defects in heterotrimeric kinesin-2 loading at the ciliary base lead to defects in the ciliary tubulin and EB1, and thus odour receptors on the ciliary shafts. A) Changes in intensity of α-Tubulin84B::GFP (i), EB1::GFP (ii), *endo*Cep290::GFP (iii), *endo*CEP135::GFP (iv), KLP68D::YFP in the ciliary shaft (v), and KLP68D::YFP at the ciliary base (vi) of basiconic olfactory cilia of 1-2-day-old control flies and flies with conditional simultaneous knockdown of γ-Tubulin23C with CNN. Representative images of different fluorescently tagged ciliary markers and their corresponding quantifications are presented in the left and right, respectively. B) Changes in *endo*ORCO::GFP intensity in the basiconic olfactory cilia of 10-day-old control flies and flies with conditional simultaneous knockdown of γ-Tubulin23C with PLP and CNN with PLP. Representative images (i) and respective quantifications (ii). Time-dependent changes in the odour repulsion behaviours of 10-day-old control and conditional knockdown flies (iii). Each bar corresponds to a total of ≥60 flies measured in sets of 7-10 animals each. C) Scheme depicting the mechanism of cilia maintenance by the PCM at ciliary base. The interactive network of γ-Tubulin23C, CNN, and PLP (shown in blue) at the ciliary base is crucial for regulating the cargo loading on kinesin-2 motor responsible for cargo entry and their anterograde transport into the cilia. Removal of PCM components hampers kinesin-2 cargo loading at the ciliary base and transport into the ciliary shaft leading to defects in localisation of structural units (e.g., Tubulin, EB1) and functional units (e.g. Receptors) into the cilia, thereby causing loss of structural and functional integrity. Scale bars on confocal micrographs in A (i-v) and B are 10 µm each while those in A (vi) are 2.5 µm.

**Figure 7:**
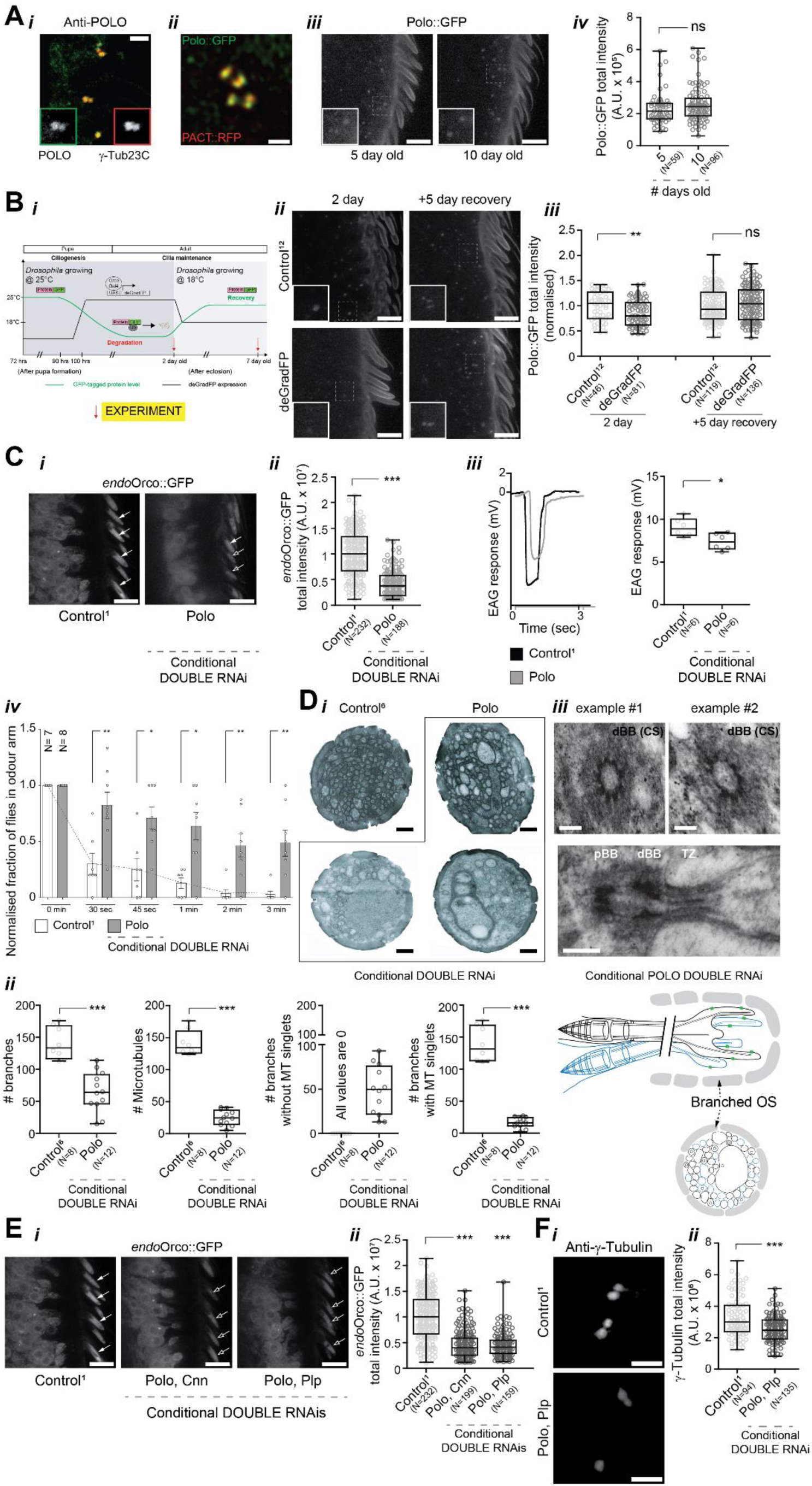
POLO kinase and PCM structural proteins work synergistically at the ciliary base for maintenance of the structure and function of adult basiconic olfactory cilia. A) Representative images of POLO localisation with γ-Tubulin in basiconic olfactory cilia of control adult flies (i). Representative Airyscan 2.0 images describe the localisation of POLO with PACT used as centriole wall marker in olfactory cilia (ii). Time-dependent changes in *endo*POLO::GFP intensity at the ciliary base in basiconic olfactory cilia of 5-day and 10-day-old adult flies. Representative images (iii) and respective quantifications (iv). B) Scheme of the approach and timeline of the conditional degradation experiments (i). Time-dependent changes in *endo*POLO::GFP intensity at 25° upon conditional degradation (under Gal4*^Orco^* driver) for 2 days and recovery (at 18°C) for 5 days post-degradation. Representative images (ii) and respective quantifications (iii). C) Changes in *endo*ORCO::GFP intensity in basiconic olfactory cilia of control and conditional POLO DOUBLE RNAi knockdown flies. Representative images (i) and respective quantifications (ii). EAG response measured from basiconic region of third antennal segment of 10-day-old control and conditional knockdown flies (iii). EAG traces (left) and respective quantifications (right) for flies with relevant genotypes. Time-dependent changes in the odour repulsion behaviours of control and conditional knockdown flies (iv). Each bar corresponds to a total of ≥60 flies measured in sets of 7-10 animals each. D) Representative electron micrographs of individual cross sections of ciliary OS (i) and branch quantification (ii) in control and conditional knockdown flies. Representative electron micrographs of individual cross sections of BBs in OSNs (iii, top) and longitudinal section (iii, middle) showing the pBB, dBB, and TZ. Scheme representing OSNs innervating a basiconic sensilla (iii, below) showing various features of the olfactory cilia namely pBB and dBB, TZ, and defective branching in the OS upon POLO removal. E) Changes in *endo*ORCO::GFP intensity in control and flies with conditional simultaneous knockdown of POLO with CNN and POLO with PLP. Representative images (i) and respective quantifications (ii). F) Changes in γ-Tubulin intensity in basiconic olfactory cilia in control and conditional knockdown flies. Representative images (i) and respective quantifications (ii). Scale bars on micrographs in A (i) and F are 2.5 µm, those in A (iii), B, C, and E are 10 µm, those in A (ii), D (i), and D (iii, middle) are 500 nm, while those in D (iii, top) are 100 nm.

**Figure 8:**
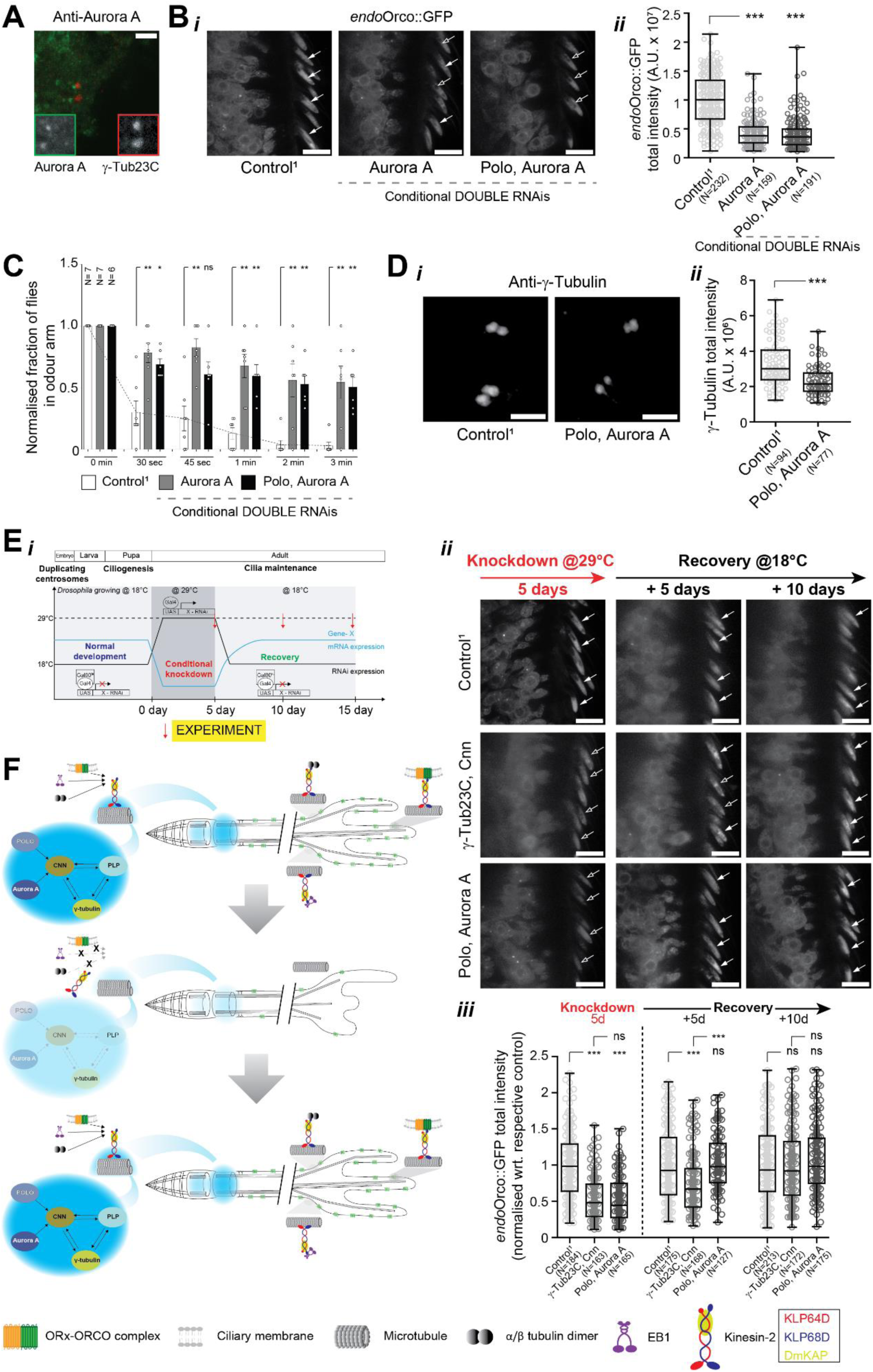
POLO and Aurora A work synergistically with PCM structural proteins at the ciliary base for maintenance of structure and function and regrowth of adult basiconic olfactory cilia. A) Representative images of Aurora A localisation with γ-Tubulin in basiconic olfactory cilia of control adult flies. B) Changes in *endo*ORCO::GFP intensity in basiconic olfactory cilia of control flies and flies with conditional DOUBLE RNAi knockdown of Aurora A and conditional simultaneous knockdown of POLO with Aurora A. Representative images (i) and respective quantifications (ii). C) Time-dependent changes in the odour repulsion behaviours of control and conditional knockdown flies. Each bar corresponds to a total of ≥60 flies measured in sets of 7-10 animals each. D) Changes in intensity of γ-Tubulin in basiconic olfactory cilia of control and conditional knockdown flies. Representative images (i) and respective quantifications (ii). E) Scheme of the approach and timeline of the conditional knockdown and recovery experiments (i). Time-dependent changes in *endo*ORCO::GFP intensity in the basiconic olfactory cilia of control and conditional knockdown flies for 5 days (29°C) and recovery (18°C) for 5 days and 10 days post- knockdown. Representative images (ii) and respective quantifications (iii). ORCO intensities were normalised wrt. control of that condition for comparison since flies were grown at different temperatures for knockdown and recovery, which could affect the overall rate of protein synthesis and turnover. F) Scheme representing proposed mechanism and regulation of cilia maintenance. The ciliary base has BBs connected by rootlet and surrounded by PCM (in blue), and a TZ. The axoneme extends into the OS with extensive branching, each membrane branch carrying MT singlets. The ciliary membrane hosts odorant receptor-ORCO complexes essential for odour sensing. Under normal homeostasis, the synergistic action of PCM proteins and regulatory kinases allows cargo loading with kinesin-2 motors at the ciliary base and transport into the cilia. Correct localisation of ciliary structural units (EB1, tubulin) and functional units (ORCO, odorant receptors) leads to maintenance of cilia structure and function. Upon conditional deregulation of the PCM, cargo loading and transport is affected leading to severe structural defects, and thus, functional defects in olfactory cilia. When the deregulation is suppressed to allow recovery of PCM, the olfactory cilia can undergo regrowth and regain its structure leading to restoration of ciliary function. Scale bars on confocal micrographs in A and D are 2.5 µm while those in B and E are 10 µm each.

Earlier studies using FRAP and *in situ* cryo-ET suggested that a pool of IFT and motor proteins resides at the ciliary base for several seconds before entry into the cilium and IFT complexes assemble at the ciliary base of *Chlamydomonas* flagella^43,44^. Also, *Drosophila* heterotrimeric kinesin-2 transports the ORCO-OR complex, α/β-tubulin, and EB1 into the fly olfactory cilia^28,45,46^. Interestingly, several key anterograde and retrograde transport proteins, including heterotrimeric kinesin-2 and cytoplasmic dynein heavy chain, were found to co-IP with CNN from the ciliary base (Supplemental Figure 5D(iii)). Therefore, to investigate whether the PCM influences the landing sites for proteins transported to the ciliary base and selectively trafficked into the cilia, we examined the localisation of KLP68D, one of the two motor subunits of heterotrimeric kinesin-2. In controls, this motor subunit gets enriched at the ciliary base, along the ciliary shaft, and at the distal end of the OS^27^. KLP68D levels are significantly reduced throughout the ciliary shaft in the combination double RNAi flies (Figure 6A(v)). Strikingly, enrichment at the BB was dramatically impaired in these conditional knockdown flies compared to controls (Figure 6A(vi)). Then, to check whether other PCM components like PLP are also involved in this ciliary base network that regulates cilia maintenance, we also generated flies expressing combinations of hairpin RNAs targeting PLP with γ-Tubulin23C or CNN simultaneously. There too, we observed a dramatic reduction (by ∼52%) of ciliary ORCO (Figure 6B(i, ii)). These combination double RNAi flies also exhibited significant defects in odour repulsion responses to external stimuli (Figure 6B(iii)).

In summary, our findings demonstrate that the PCM surrounding the BBs of long-lived olfactory cilia consists of a network of various proteins, including γ-Tubulin23C, CNN, and PLP that previously have been demonstrated to be involved in elaborating PCM around duplicating centrosomes. This network of PCM proteins is required to maintain an appropriate chemical environment to regulate motor-cargo docking from the cell body to the ciliary base prior to trafficking into the ciliary shaft, thereby influencing ciliary membrane structure, composition, function, and ciliary skeleton organisation. (summarised in Figure 6C).

### PLK1/POLO and Aurora A work together on γ-Tubulin23C, CNN, and PLP at the base of the cilia to keep the cilia shaft stable in adult flies

Given the conditional loss of PCM proteins induces change in the dynamicity of the environment around the BBs, we then aimed to investigate the mechanisms regulating these maintenance processes, specifically the organisation and activity of the ciliary base PCM. Several kinases, including PLK1, Aurora A, CDK1, NEK2, and PKA, are extensively studied to understand their regulatory roles in the PCM formation and expansion at the centrosome in cycling cells (reviewed in^47,48^). PLK1/POLO promotes the accumulation of scaffolding materials (e.g., ANA1/CEP295, CNN) at the centriole surface and recruiting γ-Tubulin ring complexes (γ-TuRC) necessary for stabilising cytoplasmic MTs around the centrosome^49,50^. Also, PLK1/POLO disappears from the centrosomes in the *Drosophila* oocytes and from centrioles at the base of *C. elegans* sensory cilia at the early stage of ciliogenesis, which precede the centrioles disappearance in these systems^19,21^. Works using hRPE-1 cells induced cilia disassembly *in vitro* model showed that PCM-1 recruits CDK1-dependent phosphorylated PLK1 to the TZ, where PLK1 works with nephronophthisis (NPHP1), HDAC6 and DVL2 to facilitate cilia disassembly^51–53^. In contrast, *C. elegans* ciliary base function during the later phase of ciliogenesis is regulated by SPD-5/CNN, γ-Tubulin and PCMD-1/PLP in a PLK1-independent process^20,21^. All these conflicting results suggest that we should first explore PLK1/POLO’s localisation at the base of these long-lived *Drosophila* olfactory cilia as localisations and roles of any centrosomal kinases in the metazoan cilia homeostasis *in vivo* are unknown.

Interestingly, we found POLO enriched at the ciliary bases primarily to the BBs (Figure 7A(i, ii)) Furthermore, POLO maintains its localisation at the ciliary base, with protein levels at the ciliary bases remaining consistent between 5-day- and 10-day-old flies (Figure 7A(iii, iv)). To determine whether POLO is dynamically replenished at the ciliary base, as observed for CEP135 and CNN (Figure 3E), an experiment was conducted (Gal4*^Orco^*-*UAS*deGradFP- *endo*POLO::GFP) involving the spatiotemporal expression of deGradFP (Figure 7B(i)). After two days of adult life, levels of endogenously expressed POLO::GFP at the olfactory cilia base were significantly reduced; upon cessation of deGradFP expression in the same cells, POLO::GFP levels returned to normal compared to controls (Figure 7B(ii, iii)). These findings indicate that POLO kinase persists at the ciliary base in the adults, and that the kinase is dynamically exchanged at the ciliary base.

Notably, ciliary ORCO was dramatically reduced in OSNs with POLO double RNAi (∼60%) compared to control (Figure 7C(i, ii)). The conditional POLO knockdown flies show reduced EAG response (7.4±1.0 mV) than control (9.1±1.1 mV) (Figure 7C(iii)) and impaired odour response behaviour compared to control flies (Figure 7C(iv)). We then found the number of branches at the OS of POLO double RNAi olfactory cilia is also significantly reduced (65±30) compared to control (140±25) (Figure 7D(i, ii)). The gross organisation of the ciliary bases (BBs and TZ) were visibly unaffected (Figure 7D(iii)). Collectively, these results demonstrate the importance of POLO kinase in maintaining cilia after ciliogenesis is completed. Then, to test the hypothesis that POLO kinase works with the PCM proteins, whose role we studied in the earlier section, we generated flies with a combination knockdown of one of the three key centrosome components with POLO. We found that OSN- and adult-specific knockdown of POLO with CNN or POLO with PLP leads to a significant reduction (∼55%) in ciliary ORCO (Figure 7E), and in these flies, the γ-Tubulin level at the ciliary base is significantly reduced (Figure 7F), suggesting that POLO kinase might be affecting the function of the PCM at the BB, which in turn is affecting the cilia homeostasis.

Note that the efficacy of the RNAi hairpins were validated using a ubiquitously active promoter (Gal4*^Tubulin^*) (Supplemental Figure 7A) and they were found to phenocopy the null mutant described previously^54^. Also, the POLO mRNA levels in antennal tissue of conditional knockdown flies were significantly reduced (by 95%, Supplemental Figure 7B). Still, the defects observed in the ciliary OS in POLO double RNAi flies were weaker when compared to PCM knockdowns (Figure 5E, 7D and Supplemental Figure 6). This made us hypothesise that POLO kinase may not be sufficient for the process of cilia maintenance that we are investigating and may require other kinases.

We decided to investigate Aurora A, another critical kinase known to promote the accumulation of scaffolding materials at the centriole surface during centrosome maturation^55^. Also, though the localisation of Aurora A is ambiguous at the ciliary base, phosphorylated Aurora A/CALK is implicated in cilia disassembly in hRPE-1 cells and *Chlamydomonas*, and ectopic expression of Aurora A leads to fast cilia disassembly^56–58^. Together with the above facts and given Aurora A’s localisation at the base of any long-lived metazoan cilia *in vivo* is unknown, we first analysed the localisation of Aurora A in the *Drosophila* adult olfactory ciliary base. Interestingly, Aurora A localises to the proximal BB and the rootlet (Figure 8A). Notably, in contrast to our conjecture based on *in vitro* works earlier reported, we found the ciliary ORCO was significantly reduced (by ∼58%) in OSNs with Aurora A double RNAi knockdown (Figure 8B(i, ii)). Furthermore, simultaneous removal of POLO with Aurora A resulted in a dramatic reduction (by ∼60%) of ciliary ORCO localisation (Figure 8B(i, ii)) and its volume (∼45%) (Supplemental Figure 7C). These combination double RNAi flies also exhibited significant defects in olfactory responses to external stimuli (Figure 8C) and reduced ciliary base localisation of γ-Tubulin (Figure 8D).

In summary, we discovered a novel combinatorial role of POLO and Aurora A at the ciliary base to maintain PCM chemistry that is required to maintain ciliary shaft structure and function *in vivo*.

### *Drosophila* olfactory cilia are dynamic and capable of regrowth *in vivo*

In *Sea urchin* and *Xenopus embryo* and *Chlamydomonas,* chemically induced shaved off flagella can regenerate^59–61^. The mouse photoreceptor OS (membrane disc), which lacks the MT-skeleton, is replenished daily^62^. Studies in *C. elegans* sensory cilia suggest IFT rates and the sensory functions decrease with age, but cilia length remains same during worms adulthood^16,63–65^. Also, vertebrate sperm flagella, *Trypanosoma* flagella, and *Drosophila* auditory cilia skeleton structure remain stable for the lifetime of these cells^13,40,66^. Yet, the regeneration ability of ectopically damaged metazoan cilia skeleton structure within tissue has just begun to be explored, likely due to limited availability of appropriate tools and techniques. Therefore, we investigated whether structurally and functionally damaged cilia could recover to their normal morphology and physiology after stopping the spatiotemporal knockdown. To do this, we used the temperature-sensitive Gal4-*UAS*-*Tub*Gal80^ts^ system (Figure 8E(i)). 5 days of knockdown of γ-Tubulin23C with CNN or POLO with Aurora A leads to reduction of ciliary ORCO levels (by ∼45-48%, Figure 8E(ii, iii)) and caused odour repulsion defects (Supplemental Figure 8A(i)) consistent with previously described results on 10 day knockdown (Figure 5 and 8), after which we shifted the flies to 18°C to promote recovery. Over time, ORCO levels in the ciliary shaft increased, and by 10 days after stopping knockdown, ciliary ORCO levels were similar to controls (Figure 8E (ii, iii)). The adult flies with olfactory cilia also showed normal odour repulsion behaviour indistinguishable from controls (Supplemental Figure 8A(ii, iii)). Moreover, the defect caused by POLO and Aurora A removal fully recovered by 5 days, suggesting the ciliary shaft regrew faster when kinase removal was suppressed than when PCM structural proteins were restored (Figure 8E (ii, iii)). This is also reflected in the slower olfactory behavioural recovery of γ-Tubulin23C with CNN knockdown flies compared to POLO with Aurora A knockdown flies (Supplemental Figure 8A (ii, iii)). Then, in a parallel experiment with a 10-day knockdown (Supplemental Figure 8B(i)), we observed a more pronounced reduction in ORCO levels (by ∼62%) linked to a significant odour repulsion defect, as expected from the extended knockdown period (Supplemental Figure 8B(ii, iii) and C(i)). Surprisingly, recovery was also seen in this group of cilia and flies, but at a slower rate (Supplemental Figure 8B(ii, iii) and C(ii, iii)). This slower recovery may result from the more substantial initial knockdown or the increased age of the flies, which could slow recovery due to age-related decline.

In summary, these results indicate that olfactory cilia are dynamic and capable of regrowth *in vivo*. Recovery from damage in the skeleton structures and membrane compositions is directly associated with ciliary functionality and the animal’s pathophysiology in adulthood.

## Discussion

### Adult *Drosophila* OSNs: a powerful, physiologically relevant *in vivo* sub-cellular compartment/organelles homeostasis investigation platform

Here, we showed that adult *Drosophila* OSNs maintain stable ciliary structure and function over 5- to 10-day period. Ciliary structural (protein compositions, localisations and electron microscopy) and functional (electrophysiological response and behavioural assay) analyses of cilia suggest active, cell-autonomous maintenance of structure and function during young adulthood (Figure 1-3, Supplemental Figure 1-3). The consistent ultrastructural features, including BB and TZ architecture, and stable numbers of OS branches, suggest that ciliary homeostasis is energetically demanding and requires ongoing cellular active regulation. We then, for the first time, show that the BB (structural proteins and kinases) serves as a central coordinator for BB-dendritic MT anchoring, loading of Kinesin-2 to the ciliary base, thus motor/cargo entry to the TZ and ciliary shaft, all of which are essential for maintaining ciliary delicate shape and vital function (Figure 4-8, Supplemental Figure 4-8).

The studies in *Chlamydomonas* flagella and *C. elegans* sensory cilia revealed that IFTs undergo continuous turnover^11,12,14,15^. *Chlamydomonas* and mouse culture cell lines flagella/cilia length is tightly regulated through motor-cargo transports mechanisms adjusting the balance between assembly and disassembly^11,67^. Our study extends these principles of ciliary shaft homeostasis to a complex, sensory cilium within a multicellular organism, demonstrating that ciliary dynamic maintenance is conserved in metazoans *in vivo*. For the first time, we showed that the resorbed metazoan cilia under deregulated condition *in vivo*, once deregulation process is suppressed, can regrow to its full functionality cell autonomously (Figure 8E, Supplemental Figure 8).

In summary, these results demonstrate that adult *Drosophila* OSNs are robust, genetically tractable, and serve as a physiologically relevant *in vivo* model for investigating the mechanisms of ciliary homeostasis (Figure 1-8, Supplemental Figure 1-8). This system is also applicable to studying various phenomena in other compartments/organelles, including mitochondria, the Golgi apparatus, lysosomes, autophagosomes, P-bodies, as well as related aspects of cellular homeostasis.

Importantly, although this study offers many new details on the structural and functional stability of the ciliary base and shafts, several limitations remain. For instance, here we did not aim to associate our study with the impact of ageing on cilia homeostasis. Future longitudinal studies that track individual sensilla over the full 60-day *Drosophila* adult lifespan or compare very old flies (50-60 days) with young and middle-aged adults, could determine whether homeostatic mechanisms eventually fail or adapt.

### PCM at ciliary base forms a dynamic meshwork similar to duplicating centrosome but with altered molecular composition and organisation

We found that the BBs of *Drosophila* olfactory cilia contain a specialised, dynamically maintained subset of centrosomal proteins. CEP135/BLD10, γ-Tubulin, CNN, and PLP remain at the BBs for at least 10 days in adults. In contrast, SAS6, ANA2, and ASL are excluded, indicating a distinct molecular composition from that of duplicating centrosomes. The absence of these proteins suggests that olfactory BBs might not be able to template pro-centriole formation, but they are functional MTOCs as they retain γ-Tubulin and other PCM structural proteins. The organisation of retained PCM proteins around BBs differs from that in centrosomes in cycling cells, showing partial overlap among γ-Tubulin, CNN, and PLP and an altered radial distribution (Figure 2, Supplemental Figure 2)^31,32^. Though *Chlamydomonas* flagella base is considered to be ciliary transport assembly site^43,44^, the BB (i.e., modified centrosome) proteins’ dynamicity was never investigated. Using conditional degradation and recovery experiments, i.e., pulse-chase-pulse system, we showed that at least a few centriole and PCM proteins persisting at the ciliary base undergo active turnover (Figure 3, Supplemental Figure 3). These first-of-their-kind findings enhance our understanding of centrosome modifications and underscore the importance of dynamic regulation of PCM in ciliary homeostasis in a mature tissue. Further work is needed to determine how many other centriole and PCM components are dynamically regulated, and further analyses on how conserved centrosome modification at various ciliary bases with centrioles across diverse cell types and species are needed.

### Kinases are regulators of ciliary base PCM organisation, dynamicity and function

We, for the first time in any ciliated cell model system, showed that γ-Tubulin23C, CNN, and PLP are required for the maintenance of olfactory cilia function in adult *Drosophila*, independent of their roles in centriole duplication during development (Figure 4, Supplemental Figure 4). We then showed that γ-Tubulin23C, CNN, and PLP co-IP from ciliary base-enriched fractions, and that the combinatorial knockdown of PCM proteins produces synergistic ciliary defects, supporting their interdependence in maintaining BB organisation and function (Figure 5, 6, and Supplemental Figure 5, 6). To avoid confounding effects on ciliary function and animal behaviour, we perturbed only selected OSNs by choosing a conditional RNAi driver (Gal4*^Orco^*) with mild expression that led to ≥60% knockdown in the antenna using double RNAis against a single gene. Note that this estimation of mRNA reduction is an underestimation as knockdown was conducted only in ∼70% of the OSNs, but quantitative RNA analysis was done using second and third antennal tissues, which comprise cells more than those OSNs targeted by Gal4*^Orco^*. Importantly, e.g. adult-specific simultaneous knockdown of γ-Tubulin23C with CNN in OSNs using double RNAis caused a dramatic reduction of enriched γ-Tubulin at the ciliary base, ∼80% reduction in ciliary branches, and ∼60% reduction in ORCO (Figure 4-6 and Supplemental Figure 4-6). Ciliary maintenance defects in OSNs innervating all three classes of sensilla (basiconic, trichoidic, and coeloconic) in γ-Tubulin23C double RNAi flies suggest that mechanisms proposed for basiconic olfactory cilia could also apply to other types of *Drosophila* olfactory cilia (Supplemental Figure 5A, B). Still, in principle, a more pronounced knockdown, thus a pronounced ciliary defect, can be achieved using a stronger Gal4 driver (Gal4*^ChAT19b^*), a tool we used in an earlier study, but that is not restricted to OSNs and broadly expresses across all cholinergic neurons, which include all Type-I sensory neurons^40^.

Then we showed *Drosophila* POLO/PLK1 and Aurora A localise at the ciliary base. While POLO localises at the walls of both BBs, Aurora A enriches at the proximal BB and rootlet. These localisation patterns differ from those reported in the ciliary base of hRPE-1 cells^51,57,58^. At the OSN ciliary bases, we showed for the first time that kinases undergo dynamic turnover like centrosomal proteins (Figure 7B). Our data suggest POLO and Aurora A act cooperatively and function with or partially upstream of PCM network of proteins (CNN, γ-Tubulin23C, and PLP). They regulate PCM organisation to maintain ciliary structure and function (Figure 7, 8 and Supplemental Figure 7, 8). Earlier works using a ciliary disassembly model suggested that these kinases’ activity at the ciliary TZ and BB leads to cilia disassembly ^52,53,56–58^. In contrast, our findings reveal these conserved centrosomal kinases are actively required to preserve the olfactory cilia structure and function. They also expand the functional repertoire of POLO and Aurora A beyond mitotic cell division. Our data show that structural PCM proteins and kinases at the BB are not static remnants of ciliogenesis but are actively required for ongoing ciliary maintenance (Figure 3, 6-8, Supplemental Figure 3, 6-8).

These observations also provide an important rationale for how mutations/deregulations in components of ciliary base, including centrosome/BB (e.g., ALMS1, Pericentrin/PCNT, α- actinin/ACTN1) and TZ (e.g. NPHPs, inversion, CEP164) can generate the complex tissue-specific late-onset phenotypes observed in human ciliopathies, like Alström syndrome, Senior-Loken syndrome (SLS), Retinal Degeneration, Joubert syndrome^5,7,8^. The proposed mechanism suggests a broader relevance for sensory disorders involving compromised ciliary maintenance. Also, future research could examine whether the proposed mechanisms are conserved across various ciliates and, if so, to what extent.

### PCM proteins at ciliary base recruit ciliary motor proteins providing regeneration capacity to a metazoan cilium *in vivo*

PCM proteins, particularly γ-Tubulin23C, CNN and PLP, are critical regulators of ciliary shaft architecture and function in mature, non-dividing OSNs. We propose that PCM proteins operate through several coordinated mechanisms: (1) maintaining γ-Tubulin localisation and MTOC activity at the BB; (2) facilitating the recruitment and concentration of the primary IFT motor Kinesin-2 at the ciliary base; and (3) regulating the localisation of MT-associated proteins like EB1 within the ciliary shaft, thereby influencing tubulin dynamics and MT organisation in the cilia. The observed reduction of the ciliary OS, together with preservation of the BB core structure, i.e., 9-fold symmetric centriole-MT organisation, demonstrates a functional separation between the structural and catalytic roles of PCM proteins. This difference between ciliary assembly-promoting and MTOC-regulating PCM proteins provides a new framework for understanding ciliary development, homeostasis and dysfunction in ciliopathies. Our findings indicate that the BB functions as an active, dynamic control centre for ciliary organisation, rather than serving solely as a passive foundation, thereby advancing our understanding of ciliary homeostasis in differentiated cells *in vivo* (Figure 4-6, Supplemental Figure 4-6).

Furthermore, we have, for the first time, proposed a molecular mechanism of the regenerative potential of mature sensory cilia (Figure 8E, F and Supplemental Figure 8), suggesting new opportunities for therapeutic intervention in diseases presumed to be caused by ciliary degeneration. Also, several open questions remain. For example, though we have studied five key proteins, many other centriole and PCM components have yet to be characterised in this context. Also, here we failed to remove centrioles/BBs from the ciliary base, suggesting a hitherto unknown, but complex active process underlies the persistence of centrioles at the ciliary base. Future research should use spatial proteomics, cryo-electron microscopy of isolated ciliary bases, live-cell imaging of transport dynamics, structure-function analysis of PCM protein interactions, and systems biology to achieve a comprehensive mechanistic understanding of how ciliary base proteins coordinate homeostasis processes like maintain MT organisation, protein trafficking, membrane protein distribution, and axonemal architecture, which are required to maintain ciliary structure and function throughout the cilia, cells, and organisms’ lifespan.

## Acknowledgement

We thank Krishanu Ray, Renata Basto, Jordan Raff, and Benedicte Durand for reagents. We acknowledge Sanskruti Jagadesh and Sumiran Kasturi for help in some ubiquitous knockdown experiments, Ragini Raghu for helping in olfaction assay standardisation, Merrin Vincent for initiating tissue-specific centrosome purification and IP standardisations, and Vineet Kumar for initiating electrophysiology set-up assembly. We thank Swadhin Chandra Jana (SCJ) Lab (Organelle Biology Lab) members for reviewing the manuscript and providing helpful discussions on the manuscript. We thank the IGC/GIMM light and electron microscopy, and fly facilities, the NCBS Mass-Spectrometry Facility, NCBS Central Imaging & FACS Facility, and NCBS Electron Microscopy Facility for helping us with data acquisition, and the NCBS fly facility for assisting us with fly husbandry and fly imports. Minita Desai (MD), Pranjali Priya (PP), Merrin Vincent and P Okenve-Ramos (POR, PTDC/BIA-BID/32225/2017) are supported by the NCBS-Tata Institute for Fundamental Research (TIFR), CSIR (Council of Scientific & Industrial Research, India) and FCT (Fundação Portuguesa para a Ciência e Tecnologia, Portugal) Fellowships/Grants/Contracts. MD received travel support from Infosys (NCBS-TIFR) and financial grants from the Anusandhan National Research Foundation (ANRF), India and ASCB-EMBO travel grant, USA-Europe. PP received travel support from Infosys (NCBS-TIFR). Authors acknowledge NCBS-TIFR-DAE (an intramural grant from Tata Institute of Fundamental Research, Department of Atomic Energy (TIFR-DAE) to SCJ), the European Research Council Consolidator Grant (CoG683528 to M Bettencourt-Dias(MBD)), Department of Biotechnology, Government of India (BT/PR53868/BMS/85/255/2024 to SCJ) and Centre Franco-Indien pour la Promotion de la Recherche Avancée (CEFIPRA) (6903-1 to SCJ), Ministry of Earth Science, Government of India (MoES/PAMC/Dom/66/2023(E-14508) to SCJ).

## Authors Contributions

MD, PP, S, HD, PC, POR, AM and SCJ - experimentation and data analysis; MD, PP, POR, NS, MBD, and SCJ - conceptualisation and data curation; NS, MBD and SCJ - project administration; MBD and SCJ - resource management; MD, PP, and SCJ - wrote the original draft; MD, PP, POR, MBD and SCJ - reviewed and edited the drafts; and all authors commented on the drafts.

## References

1. Breslow, D. K. & Holland, A. J. Mechanism and Regulation of Centriole and Cilium Biogenesis. Annu. Rev. Biochem. 88, 691–724 (2019).

2. Jana, S. C. et al. Differential regulation of transition zone and centriole proteins contributes to ciliary base diversity. Nat. Cell Biol. 20, 928–941 (2018).

3. Li, L. & Ran, J. Regulation of ciliary homeostasis by intraflagellar transport-independent kinesins. Cell Death Dis. 15, 47 (2024).

4. Lacey, S. E. & Pigino, G. The intraflagellar transport cycle. Nat. Rev. Mol. Cell Biol. 26, 175–192 (2025).

5. Derderian, C., Canales, G. I. & Reiter, J. F. Seriously cilia: A tiny organelle illuminates evolution, disease, and intercellular communication. Dev. Cell 58, 1333–1349 (2023).

6. Zhu, X. Mammalian motile cilia: Structure, formation, organization, and function. Semin. Cell Dev. Biol. 175, 103651 (2025).

7. Reiter, J. F. & Leroux, M. R. Genes and molecular pathways underpinning ciliopathies. Nat. Rev. Mol. Cell Biol. 18, 533–547 (2017).

8. Sreekumar, V. & Norris, D. P. Cilia and development. Curr. Opin. Genet. Dev. 56, 15– 21 (2019).

9. Werner, S., Pimenta-Marques, A. & Bettencourt-Dias, M. Maintaining centrosomes and cilia. https://journals.biologists.com/jcs/article/130/22/3789/56439/Maintaining-centrosomes-and-cilia (2017) doi:10.1242/jcs.203505.

10. Tilney, L. G. & Gibbins, J. R. Differential effects of antimitotic agents on the stability and behavior of cytoplasmic and ciliary microtubules. Protoplasma 65, 167–179 (1968).

11. Marshall, W. F. & Rosenbaum, J. L. Intraflagellar transport balances continuous turnover of outer doublet microtubules. J. Cell Biol. 155, 405–414 (2001).

12. Hao, L. et al. Intraflagellar transport delivers tubulin isotypes to sensory cilium middle and distal segments. Nat. Cell Biol. 13, 790–798 (2011).

13. Fort, C., Bonnefoy, S., Kohl, L. & Bastin, P. Intraflagellar transport is required for the maintenance of the trypanosome flagellum composition but not its length. J. Cell Sci. 129, 3026–3041 (2016).

14. Song, L. & Dentler, W. L. Flagellar Protein Dynamics in *Chlamydomonas* *. J. Biol. Chem. 276, 29754–29763 (2001).

15. Loseva, E., Mitra, A., Groskamp, D. & Peterman, E. J. G. Intraflagellar transport of tubulin maintains steady-state axoneme integrity in C. elegans cilia. 2026.04.14.718528 Preprint at 10.64898/2026.04.14.718528 (2026).

16. Cornils, A. et al. Structural and Functional Recovery of Sensory Cilia in C. elegans IFT Mutants upon Aging. PLOS Genet. 12, e1006325 (2016).

17. Kochanski, R. S. & Borisy, G. G. Mode of centriole duplication and distribution. J. Cell Biol. 110, 1599–1605 (1990).

18. Simerly, C. et al. The paternal inheritance of the centrosome, the cell’s microtubule-organizing center, in humans, and the implications for infertility. Nat. Med. 1, 47–52 (1995).

19. Pimenta-Marques, A., et al. A mechanism for the elimination of the female gamete centrosome in Drosophila melanogaster. https://www.science.org/doi/10.1126/science.aaf4866 (2016).

20. Magescas, J., Eskinazi, S., Tran, M. V. & Feldman, J. L. Centriole-less pericentriolar material serves as a microtubule organizing center at the base of *C. elegans* sensory cilia. Curr. Biol. 31, 2410–2417.e6 (2021).

21. Garbrecht, J., Laos, T., Holzer, E., Dillinger, M. & Dammermann, A. An acentriolar centrosome at the *C. elegans* ciliary base. Curr. Biol. 31, 2418–2428.e8 (2021).

22. Fernandes-Mariano, C., Bugalhão, J. N., Santos, D. & Bettencourt-Dias, M. Centrosome biogenesis and maintenance in homeostasis and disease. Curr. Opin. Cell Biol. 94, 102485 (2025).

23. Fernández-Hernández, I., Hu, E. & Bonaguidi, M. A. Olfactory neuron turnover in adult Drosophila. 2020.11.08.371096 Preprint at 10.1101/2020.11.08.371096 (2020).

24. Laissue, P. P. & Vosshall, L. B. The Olfactory Sensory Map in Drosophila. in Brain Development in Drosophila melanogaster (ed. Technau, G. M.) 102–114 (Springer, New York, NY, 2008). doi:10.1007/978-0-387-78261-4_7.

25. Clyne, P., Grant, A., O’Connell, R. & Carlson, J. R. Odorant response of individual sensilla on theDrosophila antenna. Invert. Neurosci. 3, 127–135 (1997).

26. Chakraborty, T. S., Goswami, S. P. & Siddiqi, O. Sensory Correlates of Imaginal Conditioning in Drosophila melanogaster. J. Neurogenet. 23, 210–219 (2009).

27. Jana, S. C., Girotra, M., Ray, K. & Bettencourt, -Dias Monica. Heterotrimeric kinesin-II is necessary and sufficient to promote different stepwise assembly of morphologically distinct bipartite cilia in Drosophila antenna. Mol. Biol. Cell 22, 769–781 (2011).

28. Jana, S. C. et al. Kinesin-2 transports Orco into the olfactory cilium of Drosophila melanogaster at specific developmental stages. PLOS Genet. 17, e1009752 (2021).

29. Shanbhag, S. R., Müller, B. & Steinbrecht, R. A. Atlas of olfactory organs of *Drosophila melanogaster*. Int. J. Insect Morphol. Embryol. 28, 377–397 (1999).

30. Larsson, M. C., et al. *Or83b* Encodes a Broadly Expressed Odorant Receptor Essential for *Drosophila* Olfaction. Neuron 43, 703–714 (2004).

31. Mennella, V. et al. Subdiffraction-resolution fluorescence microscopy reveals a domain of the centrosome critical for pericentriolar material organization. Nat. Cell Biol. 14, 1159–1168 (2012).

32. Fu, J. et al. Conserved molecular interactions in centriole-to-centrosome conversion. Nat. Cell Biol. 18, 87–99 (2016).

33. Jana, S. C., Mendonça, S., Werner, S. & Bettencourt-Dias, M. Methods to Study Centrosomes and Cilia in Drosophila. in Cilia: Methods and Protocols (eds Satir, P. & Christensen, S. T.) 215–236 (Springer, New York, NY, 2016). doi:10.1007/978-1-4939-3789-9_14.

34. Jurczyk, A. et al. Pericentrin forms a complex with intraflagellar transport proteins and polycystin-2 and is required for primary cilia assembly. J. Cell Biol. 166, 637–643 (2004).

35. Muresan, V., Joshi, H. C. & Besharse, J. C. ã-Tubulin in differentiated cell types: localization in the vicinity of basal bodies in retinal photoreceptors and ciliated epithelia. J. Cell Sci. 104, 1229–1237 (1993).

36. Martinez-Campos, M., Basto, R., Baker, J., Kernan, M. & Raff, J. W. The Drosophila pericentrin-like protein is essential for cilia/flagella function, but appears to be dispensable for mitosis. J. Cell Biol. 165, 673–683 (2004).

37. Sunkel, C. E., Gomes, R., Sampaio, P., Perdigão, J. & González, C. Gamma-tubulin is required for the structure and function of the microtubule organizing centre in Drosophila neuroblasts. EMBO J. 14, 28–36 (1995).

38. Megraw, T. L., Kao, L.-R. & Kaufman, T. C. Zygotic development without functional mitotic centrosomes. Curr. Biol. 11, 116–120 (2001).

39. Tavosanis, G., Llamazares, S., Goulielmos, G. & Gonzalez, C. Essential role for γ- tubulin in the acentriolar female meiotic spindle of Drosophila. EMBO J. 16, 1809–1819 (1997).

40. Werner, S. et al. IFT88 maintains sensory function by localising signalling proteins along Drosophila cilia. Life Sci. Alliance 7, (2024).

41. Lehmann, V., Müller, H. & Lange, B. M. H. Immunoisolation of Centrosomes from Drosophila melanogaster. Curr. Protoc. Cell Biol. 29, 3.17.1–3.17.13 (2005).

42. Tovey, C. A. et al. Autoinhibition of Cnn binding to γ-TuRCs prevents ectopic microtubule nucleation and cell division defects. J. Cell Biol. 220, e202010020 (2021).

43. Wingfield, J. L. et al. IFT trains in different stages of assembly queue at the ciliary base for consecutive release into the cilium. eLife 6, e26609 (2017).

44. van den Hoek, H., et al. In situ architecture of the ciliary base reveals the stepwise assembly of intraflagellar transport trains. Science 377, 543–548 (2022).

45. Girotra, M. et al. The C-terminal tails of heterotrimeric kinesin-2 motor subunits directly bind to α-tubulin1: Possible implications for cilia-specific tubulin entry. https://onlinelibrary.wiley.com/doi/10.1111/tra.12461 (2016) 10.1111/tra.12461.

46. Agarwal, R. G. et al. EB1 surges promote ciliary outer-segment growth through periodic tubulin influxes into the Drosophila olfactory cilia. J. Cell Sci. 139, jcs263625 (2026).

47. Langlois-Lemay, L. & D’Amours, D. Moonlighting at the Poles: Non-Canonical Functions of Centrosomes. Front. Cell Dev. Biol. 10, (2022).

48. Blanco-Ameijeiras, J., Lozano-Fernández, P. & Martí, E. Centrosome maturation – in tune with the cell cycle. J. Cell Sci. 135, jcs259395 (2022).

49. Pimenta-Marques, A. et al. Ana1/CEP295 is an essential player in the centrosome maintenance program regulated by Polo kinase and the PCM. EMBO Rep. 25, 11 (2024).

50. Conduit, P. T. et al. The Centrosome-Specific Phosphorylation of Cnn by Polo/Plk1 Drives Cnn Scaffold Assembly and Centrosome Maturation. Dev. Cell 28, 659–669 (2014).

51. Seeger-Nukpezah, T., et al. The Centrosomal Kinase Plk1 Localizes to the Transition Zone of Primary Cilia and Induces Phosphorylation of Nephrocystin-1. PLOS ONE https://journals.plos.org/plosone/article?id=10.1371/journal.pone.0038838 (2012).

52. Wang, G. et al. PCM1 recruits Plk1 to the pericentriolar matrix to promote primary cilia disassembly before mitotic entry. J. Cell Sci. https://journals.biologists.com/jcs/article/126/6/1355/54295/PCM1-recruits-Plk1-to-the-pericentriolar-matrix-to (2013).

53. Lee, K. H. et al. Identification of a novel Wnt5a–CK1ε–Dvl2–Plk1-mediated primary cilia disassembly pathway. EMBO J. 31, 3104–3117 (2012).

54. Sunkel, C. E. & Glover, D. M. Polo, a mitotic mutant of Drosophila displaying abnormal spindle poles. J. Cell Sci. 89, 25–38 (1988).

55. Magnaghi-Jaulin, L., Eot-Houllier, G., Gallaud, E. & Giet, R. Aurora A Protein Kinase: To the Centrosome and Beyond. Biomolecules 9, 28 (2019).

56. Pan, J., Wang, Q. & Snell, W. J. An Aurora Kinase Is Essential for Flagellar Disassembly in Chlamydomonas. Dev. Cell 6, 445–451 (2004).

57. Pugacheva, E. N., Jablonski, S. A., Hartman, T. R., Henske, E. P. & Golemis, E. A. HEF1-Dependent Aurora A Activation Induces Disassembly of the Primary Cilium. Cell 129, 1351–1363 (2007).

58. Plotnikova, O. V. et al. Calmodulin activation of Aurora-A kinase (AURKA) is required during ciliary disassembly and in mitosis. Mol. Biol. Cell 23, 2658–2670 (2012).

59. Auclair, W. & Siegel, B. Cilia Regeneration in the Sea Urchin Embryo: Evidence for a Pool of Ciliary Proteins. Science https://www.science.org/doi/10.1126/science.154.3751.913 (1966).

60. Rosenbaum, J. L., Moulder, J. E. & Ringo, D. L. FLAGELLAR ELONGATION AND SHORTENING IN CHLAMYDOMONAS. J. Cell Biol. https://rupress.org/jcb/article/41/2/600/476/FLAGELLAR-ELONGATION-AND-SHORTENING-IN (1969).

61. Rao, V. G., Subramanianbalachandar, V. A., Magaj, M. M., Redemann, S. & Kulkarni, S. S. Mechanisms of cilia regeneration in Xenopus multiciliated epithelium in vivo. EMBO Rep. 26, 15 (2025).

62. Young, R. W. THE RENEWAL OF PHOTORECEPTOR CELL OUTER SEGMENTS. J. Cell Biol. 33, 61–72 (1967).

63. Zhang, Y. et al. The decrease of intraflagellar transport impairs sensory perception and metabolism in ageing. Nat. Commun. 12, 1789 (2021).

64. Mitra, A., Loseva, E. & Peterman, E. J. G. IFT cargo and motors associate sequentially with IFT trains to enter cilia of C. elegans. Nat. Commun. 15, 3456 (2024).

65. Judge, K. et al. Neuron-intrinsic and glial pathways regulate sensory cilia regeneration in adult C. elegans. 2026.05.15.725580 Preprint at 10.64898/2026.05.15.725580 (2026).

66. Zhao, W. et al. Outer dense fibers stabilize the axoneme to maintain sperm motility. J. Cell. Mol. Med. 22, 1755–1768 (2018).

67. Engelke, M. F. et al. Acute Inhibition of Heterotrimeric Kinesin-2 Function Reveals Mechanisms of Intraflagellar Transport in Mammalian Cilia. Curr. Biol. 29, 1137–1148.e4 (2019).

