## Supplemental Information for "Centrosome maintains the integrity of and repairs mature olfactory cilia in adult *Drosophila*"

### Materials and Methods:

##### *Drosophila* stocks and husbandry

All fly stocks used in this study are summarised in Supplemental Table 02. Fly stocks were reared on standard cornmeal media at 25°C. For ubiquitous knockdown experiments, crosses were grown at 25°C; for conditional knockdown experiments, freshly laid eggs (at 25°C) were transferred to 18°C and grown at 18°C until fly eclosion. The eclosed flies were shifted to 29°C and again shifted back to 18°C when required, for example, for rescue of GFP degradation and conditional knockdown experiments, as described in the fly genetics section.

##### Fly genetics

For ubiquitous knockdown experiments, flies with *UAS*-RNAi of a given gene were crossed to driver lines with $Gal4^{Tubulin}$ carrying *UAS*-mCD8::GFP construct to segregate RNAi carrying GFP-positive progeny from control GFP-negative progeny. Crosses were grown at 25°C and transferred every 3 days. Pupa formation index was calculated by measuring the ratio of GFP-positive pupae to GFP-negative pupae. Fly eclosion index was calculated by measuring ratio of eclosed GFP-positive flies to GFP-positive pupae.

For conditional knockdown experiments, flies with *UAS*-RNAi of a given gene were crossed to $Gal4^{Orco}$ driver lines carrying temperature-sensitive $Gal80^{ts}$ under tubulin promoter (*Tub*$Gal80^{ts}$) and Orco::GFP under its endogenous promoter (*endo*Orco::GFP). Crosses were grown at 25°C and transferred every 2 days. Eggs were grown at 18°C to repress RNAi expression due to the presence of active *Tub*$Gal80^{ts}$ and allow normal development. Freshly eclosed flies were then shifted to 29°C to induce knockdown due to inactivation of *Tub*$Gal80^{ts}$ in Orco-expressing ciliated olfactory sensory neurons (OSNs). All knockdown experiments were performed and analysed at 10 days after eclosion to allow enough time for knockdown. To rescue the conditional knockdown, the flies were shifted back to 18°C either after 5 or 10 days of knockdown (post-eclosion) in 29°C. Experiments were performed at 5 and 10 days post-shifting to 18°C. $Gal4^{Orco}$ driver lines carrying *Tub*$Gal80^{ts}$ and *endo*Orco::GFP were also crossed to $w^{1118}$ for control experiments.

For ciliary markers visualisation, flies with *UAS*-RNAi were crossed to $Gal4^{Orco}$ driver lines expressing GFP-tagged protein of interest (namely *UAS*-α-tubulin84B::GFP, *UAS*-EB1::GFP, *endo*Cep135::GFP, *endo*Cep290::GFP) and grown at 25°C. Eclosed flies were shifted to 29°C and imaged the next day. For Klp68D visualisation, flies expressing *UAS*-Klp68D::YFP under $Gal4^{ChAT19b}$ were crossed to *UAS*-RNAi and analysed as mentioned. Flies with GFP/YFP-tagged protein of interest were also crossed to $w^{1118}$ for control experiments.

For GFP-degradation experiments, flies expressing GFP-tagged protein of interest (namely *endo*CNN::GFP, *endo*POLO::GFP, *endo*Cep135::GFP, *pUbq*PACT::GFP) were crossed to Gal4$^{Orco}$ driver lines expressing *UAS*-deGradFP and grown at 25°C and transferred every 2 days. Experiments were performed at 2 days and 10 days after eclosion. To reverse the degradation, flies were shifted to 18°C at 2 days post-eclosion, and experiments were done at 5 days post-shifting to 18°C. Flies with GFP-tagged protein of interest were also crossed to $w^{1118}$ for control experiments.

**Live tissue imaging**

For visualisation of GFP-tagged proteins in the OSNs, the whole antenna from female flies was dissected and mounted in halocarbon oil, and the large basiconic sensilla-harbouring region in the third antennal segment was imaged using an Olympus FV3000 Confocal Laser Scanning Microscope. All images were taken using 60X oil-immersion (1.4 NA) objective with an optical zoom factor of 2.5X or 3.5X.

**Immunostaining**

Adult *Drosophila* heads were dissected and arranged in tissue-freezing medium with the antenna facing up in a plastic mould. The blocks were frozen in dry ice and sectioned using the Cryostat (MEV SLEE Medical). 12-15µm sections were laid on a poly-L-Lysine coated coverslip and fixed in 4% paraformaldehyde solution containing 0.1% TritonX-100 for 1.5 hours at 4°C. After blocking in 5% BSA with 0.5% TritonX-100 in PBS for 1 hour, the sections were stained using primary antibodies overnight, followed by the required washes and secondary antibody staining for 2 hours and DAPI for 15 minutes. After a final PBS wash, coverslips were mounted in 70% glycerol/Prolong Diamond. This method is a modified version of previously described protocol[1]. Immuno-stained samples were imaged using confocal (Olympus FV3000 Confocal Laser Scanning Microscope with 60X oil-immersion 1.4 NA objective and optical zoom factor of 3.5X) or super-resolution microscopes (Zeiss LSM980 Laser Scanning Microscope with Airyscan 2.0). Antibodies used in this study are summarised in Supplemental Tables 03 and 04.

**Image acquisition and analysis**

All images are captured with image acquisition parameters consistent with their respective controls for comparison of fluorescence intensity in the basiconic region of third antennal segment.

For live-tissue and fixed tissue images, intensity measurements were done in Fiji using a macro for generating sum z-projections of GFP puncta or individual sensilla projecting orthogonal from the basiconic surface and drawing ROIs covering the puncta or sensillum from the cell body till the end of the sensillar shaft. Imaris (version 9.1.2) was used to create 3D surfaces around GFP puncta or sensilla and intensities were measured within these surfaces.
For generating representative images, Fiji macro was used for making max z-projections. The images were then upscaled by increasing pixel size eightfold and converted to 8-bit images.

**Super-resolution imaging and analysis**

For super-resolution imaging, samples were collected on poly-L-lysine-coated high-precision coverslips as mentioned in the immunostaining section. For all structured illumination microscopy (SIM) Zeiss Super-Resolution Microscope (Zeiss 980 with Airyscan 2.0), an oil-immersion Plan-Apo 1.4 NA objective and two different lasers (488 nm and 560 nm) were used. All SIM images were collected and deconvolved using Zeiss Zen software (Zeiss), all images were then processed for figure making in Fiji. In addition, note that we determined the direction and the boundary of the cilia based on the background fluorescence of one of the two fluorophores in the respective raw fluorescence micrographs.

**Transmission electron microscopy and image analysis**

Antenna were dissected, fixed, processed for chemical fixation, mounted and polymerised in resin following the published method[1,2]. Serial thin sections (60-80 nm) were cut using RMC ultramicrotome, collected on formvar-coated copper slot grids and stained with 2% uranyl acetate and Reynolds lead citrate. Samples were examined and photographed at 120 kV using a Tecnai 12 electron and Talos F200 G2 Transmission electron microscopes. Finally, the images were processed in Fiji. Total number of branches and microtubule singlets were quantified in Fiji using TEM cross-sectional images of outer segments with a diameter of 2 µm to ensure consistency in the region analysed. Branches with and without microtubule singlets were also counted to compare the defects in branching pattern with respective controls. ≥8 ciliary shafts across n=3 samples were analysed.

**Odour repulsion (T-maze) assay**

Olfaction assay was performed to study fly response to repulsive odour, namely benzaldehyde. This is a modified version of the T-maze assay described previously[3]. 7-10 flies were transferred to T-mazes post-cold anaesthesia and kept for recovery for around 45 minutes. The lid covering the control arm was supplied with paraffin oil soaked Whatman filter paper and the lid covering the odour arm was supplied with $10^{-3}$ dilution of benzaldehyde (in paraffin oil) soaked Whatman filter paper. Standardisation of odour concentration was done using 2-day-old $w^{1118}$ flies. The fly behaviour was recorded for 15 min and the number of flies in the odour arm was counted every 15 sec until 15 min after odour exposure (t=0). Fraction of flies in odour arm at a given timepoint was normalised using the formula (%flies in odour arm at time t) ÷ (% flies in odour arm at time t=0) and plotted with control for comparison of repulsion behaviour over time. ≥60 flies of each genotype were measured in sets of 7-10 animals each. All sets done across different days were complemented with two sets of $w^{1118}$ - one at the beginning and one at the end of the assay as a control for the experimental setup and to account for any variability in odour concentration.

**Negative geotaxis assay**

Negative geotaxis assays were performed as described previously[4]. 7-10 flies were transferred to 100mL measuring cylinders post-cold anaesthesia and kept for recovery for around 45 minutes. The cylinder was tapped down to bring all the flies to the bottom. The walking up behaviour was recorded post-tapping for 1 min. Total 3 recordings of 1 min each were taken with 3 min recovery in between each tapping. The number of flies above the 50 mL (half-height) mark was counted every 10 sec after tapping until 1 min. %flies above 50 mL mark was plotted with control for comparison of negative geotaxis behaviour at 20 sec after tapping. ≥60 flies of each genotype were measured in sets of 7-10 animals each.

**Electrophysiology / Electroantennogram (EAG) recording**

Electrophysiological recordings were performed using modified protocol described in previous studies[2,5]. EAG responses were recorded using in-house prepared Ag/AgCl electrodes (in a NaCl-filled glass capillary). The ground electrode was placed into the head capsule, and the recording electrode was placed in the large basiconic sensilla harbouring region on the third antennal segment. For each recording, odour puff carrying $10^{-4}$ ethyl acetate diluted in paraffin oil was delivered close to the position of the recording electrode for 500 msec using in-house LabView program controlling olfactometer. The response traces were captured using LabView software and smoothened using a 20-point moving average. Voltage peaks were measured after zero-correction in Microsoft Excel and an average of 3 recordings from each fly was plotted in GraphPad Prism. At least 6 different flies of a given genotype were analysed.

**Quantitative RNA level analysis of *Drosophila* tissues**

For ubiquitous knockdown experiments, depending on the life cycle stage at which lethality was observed, either L3 larvae, pupae (removed from pupal case), or 2-day-old adult fly (wings removed) were frozen and stored in -80°C. Whole body tissue was processed for RNA extraction. For conditional knockdown experiments, whole antennae from ~200 flies (10-day-old) were used for RNA extraction. Heads were dissected and stored in -80°C. At a time, 100 heads were taken in 1.5 mL microcentrifuge tube and plunge-frozen in liquid $N_2$ for ~30 seconds and vigorously hit against the walls of the plastic beaker to dislodge the antennae from the heads. The heads without antennae were removed from the tubes and collected separately to use as control. The antennae collected in the tube were processed further for RNA extraction.

The tissues were lysed mechanically using tissue homogeniser in liquid $N_2$ and used for RNA extraction using the HiPurA® Total RNA Miniprep Purification Kit (MB602 HiMedia) protocol. RevertAid First Strand cDNA Synthesis Kit (#K1622 ThermoFisher Scientific) was used for cDNA synthesis from isolated total RNA and a diluted aliquot of 10 ng/uL was made for using as template in quantitative PCR. Primers (summarised in Supplemental table 05) for qPCR were designed using Primer-BLAST (NCBI). The primer sequence was chosen such that primers for a given transcript bind to all possible (known) isoforms and at least one of the primers is specific to an exon-exon junction. Maxima SYBR Green/ROX qPCR Master Mix (2X) (#K0221 ThermoFisher Scientific) was used for setting up qPCR using 10 ng template cDNA. GAPDH2 and eIF1A were used as housekeeping controls. This protocol is similar to the one reported previously[6,7].

**Preparation of Tissue extract, Western blotting, and Centrosome purification**

Tissues were mechanically crushed using tissue homogeniser in liquid $N_2$, followed by lysis in the lysis buffer: 50mM Tris HCl (pH 8), 250mM NaCl, 2% NP-40, 0.5% SDS, 0.5% Na-Deoxycholate, 0.1% digitonin, 1mM DTT supplied with EDTA-free Protease inhibitor (Puregene PG-122), Phosphatase inhibitor (Sigma-Aldrich P0044) and 200mM PMSF (Roche 11359061001). Lysates were spun 4 times at 15,000 RPM for 5 mins each at 4°C in Eppendorf Centrifuge 5425R and the pellet was crushed between each spin. Supernatant was collected after a final 1-hour spin at 15,000 RPM at 4°C. 1x Laemmli buffer is added to the supernatant and boiled at 95°C for 5 mins. The samples were run on SDS-PAGE followed by standard western blotting procedures. Blocking was done in TBS supplemented with 5% BSA and 0.1% Tween-20. Primary and secondary antibody (listed in Supplemental tables 03 and 04) incubations were performed in TBS supplemented with 5% BSA and 0.05% Tween-20, while washes were performed in TBS-T (0.1% TritonX-100 in TBS).

Centrosomes were purified from Embryonic and Head extracts using a modified version of a Centrosome isolation protocol[8]. Tissue extract containing 50% sucrose was layered on top of a sucrose cushion consisting of 55% and 70% sucrose. The tubes were spun at 100,000g for 1.5 hours at 4°C in Tabletop Ultracentrifuge BC Optima Max-XP (Beckman Coulter). 9-10 fractions were collected from the bottom of the tube by poking a hole using a hot needle.

**Immunoprecipitation**

Supernatant volume collected after lysis of 200 tissues (embryos or heads) was made up to 500µL and incubated with pre-equilibrated 5µL of GFP-Trap Beads (ChromoTek GFP-Trap® Magnetic Particles M-270) for 16 hours at 4°C in an end-over-end rotator. Beads were washed 4 times in lysis buffer, resuspended in 1x Laemmli buffer and prepared for analysis by SDS-PAGE and western blotting.

**Mass Spectrometry**

The samples were run on a polyacrylamide gel till they reached 1.5 cm in the resolving gel. Gel pieces were destained with a 300 mM TEAB + 50% Acetonitrile solution, then dehydrated and incubated overnight with Trypsin (Promega). Formic acid was added to stop the reaction. Supernatant containing the peptides was transferred to a new Lobind tube and the gel pieces were incubated with extraction buffer (0.1% Formic acid in 100% ACN). Samples were dehydrated using a vacuum concentrator (SpeedVac). Dried samples were reconstituted in 20µL 0.1% Formic acid and injected with appropriate total peptide into the LC column. Samples were run in Orbitrap Fusion and Orbitrap 480 (ThermoFisher Scientific).

Mass spectra from the centrosome fraction proteome and co-immunoprecipitation (IP) proteome were analysed using DIA-NN (v1.8.1) and Proteome Discoverer (Thermo Scientific), respectively. Spectra were searched against the Uniprot *Drosophila melanogaster* FASTA database with SequestHT. N-terminal methionine excision and C-carbamidomethylation were specified as modifications. The maximum number of missed cleavages was set to one. Peptide confidence thresholds were set at 1% false discovery rate (FDR) for fraction samples and 2% FDR for IP samples. When applicable, the Match Between Runs feature was enabled for spectral analysis.

**Statistical analysis and plotting the graphs**

The All-range box plots indicate the extent of the second quartile distribution with the bisecting median bar and the whiskers indicate the data spread. The dots indicate the individual data points. The bar plots indicate mean of the dataset and error bars represent standard error of the mean.

All graphs were plotted using GraphPad Prism. Statistical testing was done using Mann-Whitney U (unpaired t-test) without assuming Gaussian distribution. Following nomenclature is used throughout the article: ns- $P > 0.05$ not significant; * - $P \leq 0.05$; ** - $P \leq 0.01$; *** - $P \leq 0.001$.

#### Supplemental Tables:

**Supplemental Table 01: Centrosomal proteins and their regulatory kinases homologues in various species (Human, *Chlamydomonas*, *C. elegans*, *Drosophila* and *Drosophila* CG number of candidate genes).**

| **Human** | ***Chlamydomonas*** | ***C. elegans*** | ***Drosophila*** | **CG number** |
|---|---|---|---|---|
| TUBG1 | γ-Tubulin | TBG-1 | γ-Tubulin 23C | CG3157 |
| TUBG1 | γ-Tubulin | TBG-1 | γ-Tubulin 37C | CG17566 |
| NEDD1 | | - | Grip71 | CG10346 |
| Centrosomin (CDK5RAP2) | | SPD-5 | CNN | CG4832 |
| Pericentrin (PCNT) | | PCMD-1 | PLP | CG33957 |
| CEP152 | | - | ASL | CG2919 |
| CEP192 | | SPD-2 | SPD2 | CG17286 |
| CPAP | | SAS-4 | SAS4 | CG10061 |
| CEP135 | Bld10 | BLD10 | BLD10 | CG17081 |
| CEP295 | | - | ANA1 | CG6631 |
| CROCC | | CHE-10 | Root | CG6129 |
| CCP110 | | CP110 | CP110 | CG14617 |
| TACC3 | | TAC-1 | TACC | CG9765 |
| STIL | | SAS-5 | ANA2 | CG8262 |
| SAS-6 | Bld12 / Sas6 | SAS-6 | SAS6 | CG15524 |
| Aurora kinase A (AURKA) | CALK | AIR-1 | AurA | CG3068 |
| Aurora kinase B (AURKB) | | AIR-2 | AurB | CG6620 |
| CDK1 | CDKA1 | CDK-1 | CDK1 | CG5363 |
| PLK4 | | ZYG-1 | SAK | CG7186 |
| PLK1 | | PLK-1 | POLO | CG12306 |

**Supplemental Table 02 (A): Fly stocks used in this study**

BDSC - Bloomington *Drosophila* Stock Center

VDRC - Vienna *Drosophila* Resource Center

| **Fly stock** | **Source and ID #** |
|---|---|
| $w^{1118}$ ; + ; + | From NCBS Fly Facility |
| *UAS*-mCherry RNAi | BDSC #35785 |
| *UAS*-Polo RNAi | BDSC #36093 |
| *UAS*-Polo RNAi | BDSC #33042 |
| *UAS*-Plp RNAi | BDSC #65231 |
| *UAS*-Cnn RNAi | BDSC #35761 |
| *UAS*-Cnn RNAi | VDRC #110415/KK |
| *UAS*-**γ**-Tubulin23C RNAi | BDSC #42799 |
| *UAS*-**γ**-Tubulin23C RNAi | VDRC #107572/KK |
| *UAS*-**γ**-Tubulin37C RNAi | VDRC #109921/KK |
| *UAS*-Aurora A RNAi | BDSC #41600 |
| *UAS*-Aurora A RNAi | BDSC #35763 |
| *UAS*-**γ**-Tubulin23C RNAi; *UAS*-**γ**-Tubulin23C RNAi | This study (generated by crossing VDRC #107572/KK and BDSC #42799) |
| *UAS*-Polo RNAi; *UAS*-Polo RNAi | This study (generated by crossing BDSC #36093 and BDSC #33042) |
| *UAS*-Cnn RNAi; *UAS*-Cnn RNAi/TM6BTb | This study (generated by crossing VDRC #110415/KK and BDSC #35761) |

| | |
|---|---|
| *UAS*-Cnn RNAi; *UAS*-**γ**-Tubulin23C RNAi | This study (generated by crossing VDRC #110415/KK and BDSC #42799) |
| *UAS*-Plp RNAi; *UAS*-**γ**-Tubulin23C RNAi | This study (generated by crossing BDSC #65231 and BDSC #42799) |
| *UAS*-Plp RNAi; *UAS*-Cnn RNAi/MKRS | This study (generated by crossing BDSC #65231 and BDSC #35761) |
| *UAS*-Plp RNAi; *UAS*-Polo RNAi/TM6BTb | This study (generated by crossing BDSC #65231 and BDSC #33042) |
| *UAS*-Polo RNAi; *UAS*-Cnn RNAi/TM6BTb | This study (generated by crossing BDSC #36093 and BDSC #35761) |
| *UAS*-Aurora A RNAi; *UAS*-Aurora A RNAi | This study (generated by crossing BDSC #35763 and BDSC #41600) |
| *UAS*-Polo RNAi; *UAS*-Aurora A RNAi | This study (generated by crossing BDSC #36093 and BDSC #41600) |
| *Orco*Gal4*endo*OrcoGFP/cyo; *Tub*Gal80$^{ts}$ | - |
| *Ubq* α-Tubulin84B::GFP; + ; + | - |
| *ChAT*Gal4*UAS*-GFP; *Tub*Gal80$^{ts}$ | - |
| *ChAT*Gal4; *UAS*-mCD8::GFP*Tub*Gal4/TM6BTb | - |
| *Orco*Gal4; TM2/TM6BTb | BDSC #26818 |
| *Orco*Gal4; *UAS*-α-Tubulin84B::GFP | This study (generated by crossing *UAS*-α-Tubulin84B::GFP line received from Krishanu Ray and BDSC #26818) |
| *Orco*Gal4; *endo*Cep135::GFP | This study (generated by crossing BDSC #26818 and BDSC #60183) |

| | |
|---|---|
| *Orco*Gal4; *endo*Cep290::GFP | This study (generated by crossing BDSC #26818 and *endo*Cep290::GFP received from Bénédicte Durand) |
| *Orco*Gal4; *UAS*-EB1::GFP | This study (*UAS*-EB1::GFP line received from Krishanu Ray crossed with BDSC #26818) |
| *UAS*-Klp68D::YFP*ChAT*Gal4/cyo | Gift from Krishanu Ray |
| *UAS*-deGradFP (II) | BDSC #58740 |
| *Orco*Gal4 (III) | BDSC #23292 |
| *UAS*-deGradFP; *Orco*Gal4 | This study (generated by crossing BDSC #58740 and BDSC #23292) |
| *pUbq*PACT::GFP; MKRS/TM6BTb | - |
| *endo*POLO::GFP | BDSC #84275 |
| *endo*CNN::GFP | BDSC #60266 |
| *endo*Cep135::GFP | BDSC #60183 |

**Supplemental Table 02 (B): Control flies used in this study**

| **Fly genotype** | **Name used in this study** |
|---|---|
| *w*; *Orco*Gal4*endo*Orco::GFP/+ ; *Tub*Gal80$^{ts}$/+ | Control[1] |
| *endo*Cep135::GFP/+ | Control[2] |
| *endo*CNN::GFP/+ | Control[3] |
| *pUbq*PACT::GFP/ +; TM6BTb/+ | Control[4] |
| *w; ChAT*Gal4/+ ; *UAS*-mCD8::GFP*Tub*Gal4/*UAS*-mCherryRNAi | Control[5] |
| $w^{1118}$ ; + ; + | Control[6] |
| *Orco*Gal4/+; *UAS*-α-Tubulin84B::GFP/+ | Control[7] |
| *Orco*Gal4/+; *UAS*-EB1::GFP/+ | Control[8] |
| *Orco*Gal4/+; *endo*Cep290::GFP/+ | Control[9] |
| *Orco*Gal4/+; *endo*Cep135::GFP/+ | Control[10] |
| *UAS*-Klp68D::YFP*ChAT*Gal4/+ | Control[11] |
| *endo*POLO::GFP/+ ; *endo*POLO::GFP/+ | Control[12] |

**Supplemental Table 03: Primary antibodies used in this study**

| **Antibody details** | **Dilution** | **Source** | **Assay** |
|---|---|---|---|
| Rabbit anti-γ tubulin | 1:1000 | Sigma-Aldrich (T5192) | Western blotting |
| Mouse anti-GFP | 1:1000 | Santa Cruz Biotechnology (SC 9996) | Western blotting |
| Rabbit anti-γ-tubulin | 1:1000 | Invitrogen (MA5-46947) ThermoFisher Scientific | Immunostaining |
| Mouse anti- γ-tubulin | 1:100 | Sigma-Aldrich (T6557) | Immunostaining |
| Rabbit anti-Cep135 | 1:1000 | Gift from Timothy Megraw | Immunostaining |
| Chicken anti-PLP | 1:1000 | In Monica Bettencourt-Dias Lab | Immunostaining |
| Rabbit anti-Aurora A | 1:500 | Gift from Renata Basto/Jordan Raff | Immunostaining |
| Mouse anti-POLO | 1:50 | Gift from David Glover | Immunostaining |
| Guinea pig anti-ASL | 1:500 | Gift from Bénédicte Durand | Immunostaining |

**Supplemental Table 04: Secondary antibodies used in this study**

| **Antibody details** | **Dilution** | **Source** | **Assay** |
|---|---|---|---|
| Donkey anti-mouse HRP | 1:10000 | Invitrogen (A16017) ThermoFisher Scientific | Western blotting |
| Goat anti-rabbit HRP | 1:10000 | Invitrogen (65-6120) ThermoFisher Scientific | Western blotting |
| Donkey anti-mouse 568 | 1:1000 | Invitrogen (A10037) ThermoFisher Scientific | Immunostaining |
| Donkey anti-mouse 647 | 1:1000 | Invitrogen (A32787) ThermoFisher Scientific | Immunostaining |
| Donkey anti-rabbit 568 | 1:1000 | Invitrogen (A10042) ThermoFisher Scientific | Immunostaining |
| Donkey anti-rabbit 647 | 1:1000 | Invitrogen (A32795) ThermoFisher Scientific | Immunostaining |
| Goat anti-chicken 568 | 1:1000 | Invitrogen (A11041) ThermoFisher Scientific | Immunostaining |
| Goat anti-chicken 647 | 1:1000 | Invitrogen (A32933) ThermoFisher Scientific | Immunostaining |
| Goat anti-guinea pig 568 | 1:1000 | Invitrogen (A11075) ThermoFisher Scientific | Immunostaining |

**Supplemental Table 05: Primers for qPCR used in this study**

| ***Drosophila* CG number of the gene** | **Target (Forward / Reverse primer)** | **Primer sequence** | **Primer length** | **Primer $T_m$ (°C)** | **Amplicon size** |
|---|---|---|---|---|---|
| CG3157 | γ-Tubulin23C FP | CTCGGTCTACTCCAAGCTCT | 20 | 58.24 | 147 bp |
| | γ-Tubulin23C RP | CCATCTGCCTCACGGGTCAAT | 20 | 60.11 | |
| CG4832 | CNN FP | CATCACGAGGAATTGCAGCG | 20 | 59.97 | 100 bp |
| | CNN RP | CACCAGACTCCGCCATCTGA | 20 | 61.61 | |
| CG33957 | PLP FP | GGCAGTTGATCGGTCGCTAT | 20 | 60.25 | 190 bp |
| | PLP RP | GCAAGGGCTACCGTCTTAAA | 20 | 57.9 | |
| CG17566 | γ-Tubulin37C FP | AACAACAACCTCATCGGTCTGA | 22 | 59.9 | 110 bp |
| | γ-Tubulin37C RP | ACGCTTGTCTTGGTCTCGCA | 20 | 62.36 | |
| CG12306 | POLO FP | TATCAACGGAAAGCCGCGA | 19 | 59.78 | 106 bp |
| | POLO RP | GTCGCTGTAGTCAACCCACT | 20 | 59.68 | |
| CG8893 | GAPDH2 FP | TGCAAGCAAGCCGATAGATAA | 21 | 57.81 | 139 bp |
| | GAPDH2 RP | ATGAAGGGATCGTTGACGGC | 20 | 60.46 | |
| CG8053 | eIF1A FP | GATATACTGGTTCCCCGCGA | 20 | 59.04 | 102 bp |
| | eIF1A RP | GGCTTGTTGGCGACCAATTTT | 21 | 60.54 | |

#### Supplemental Figures:

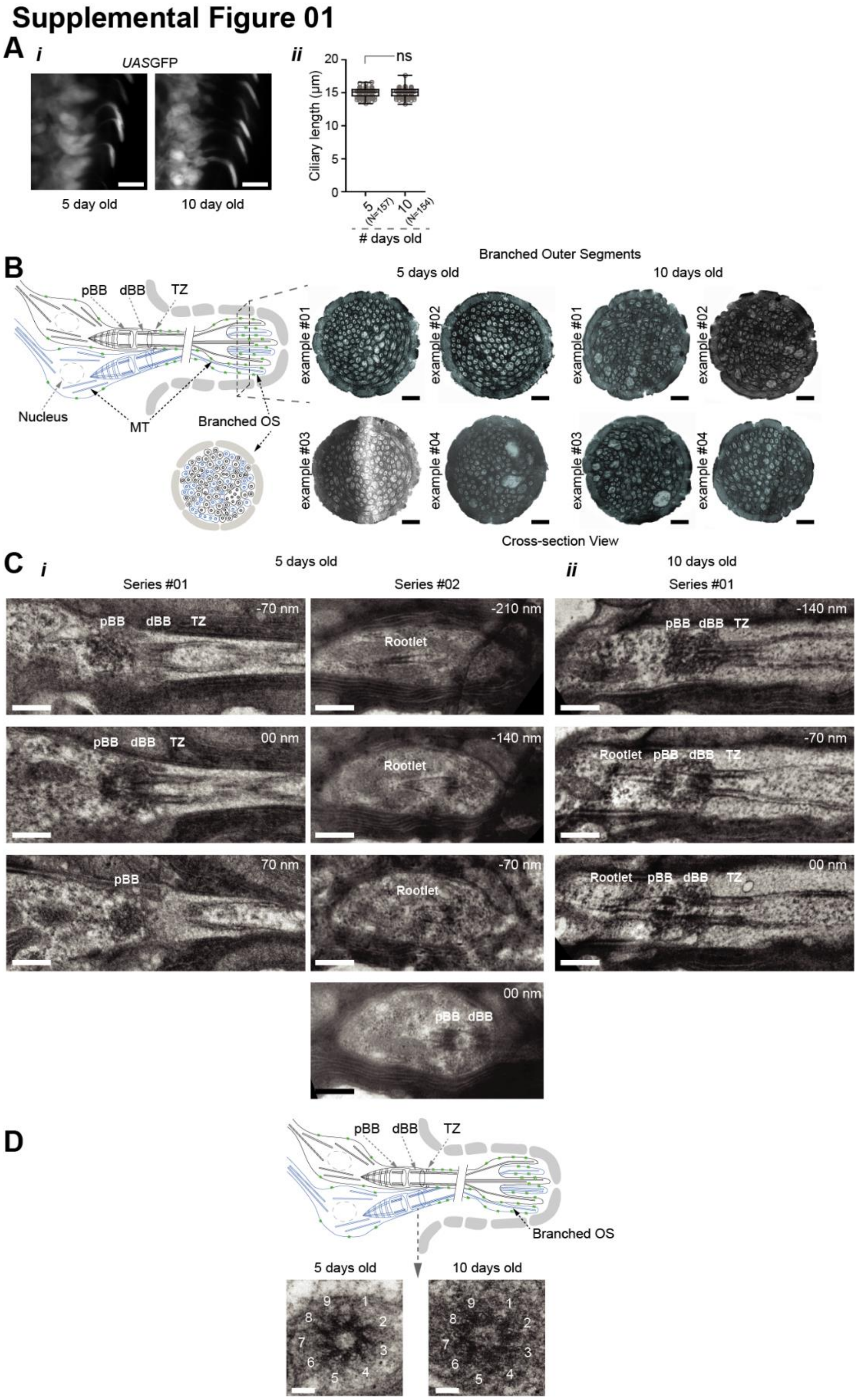

**Supplemental Figure 01: Olfactory basiconic cilia length and ultrastructural organisation are maintained between 5 and 10 day age of adult flies.**

A) Representative images (i) and ciliary length quantification (ii) of basiconic olfactory cilia in the third antennal segment of 5-day and 10-day-old adult control flies (Gal4$^{ChAT19b}$>*UAS*GFP; *UAS*-mCherryRNAi). The length of GFP fluorescence in the sensillar shaft starting from the cell body was measured to calculate ciliary length (~15 µm).

B) Representative electron micrographs of individual cross sections of branched OS from 5-day and 10-day-old control adult flies. Four different examples of each condition are shown to demonstrate the organisation of MT singlets surrounded by membrane branches quantified in Figure 1E(ii).

C) Representative electron micrographs of sets of serial longitudinal sections of BBs in OSNs from 5-day (i) and 10-day-old (ii) control adult flies.

D) Representative electron micrographs of individual cross sections of BBs in OSNs from 5-day (left) and 10-day-old (right) adult flies showing 9-fold symmetry.

Schemes in B and D represent OSNs innervating a basiconic sensilla. Various features of the olfactory cilia namely pBB and dBB, TZ, and branched OS are depicted for comparison with the micrographs.

Scale bars on confocal micrographs in A are 10 µm. For the longitudinal-section series analysis, ~70 nm serial sections were collected. All electron micrographs in B, C represent features observed in n=3 samples (the experiments were repeated independently at least twice with similar results). Scale bars on the OS cross section (B) and BB longitudinal section (C) micrographs are 500 nm each while that on BB cross section micrographs (D) are 100 nm.

### Supplemental Figure 02

A

B

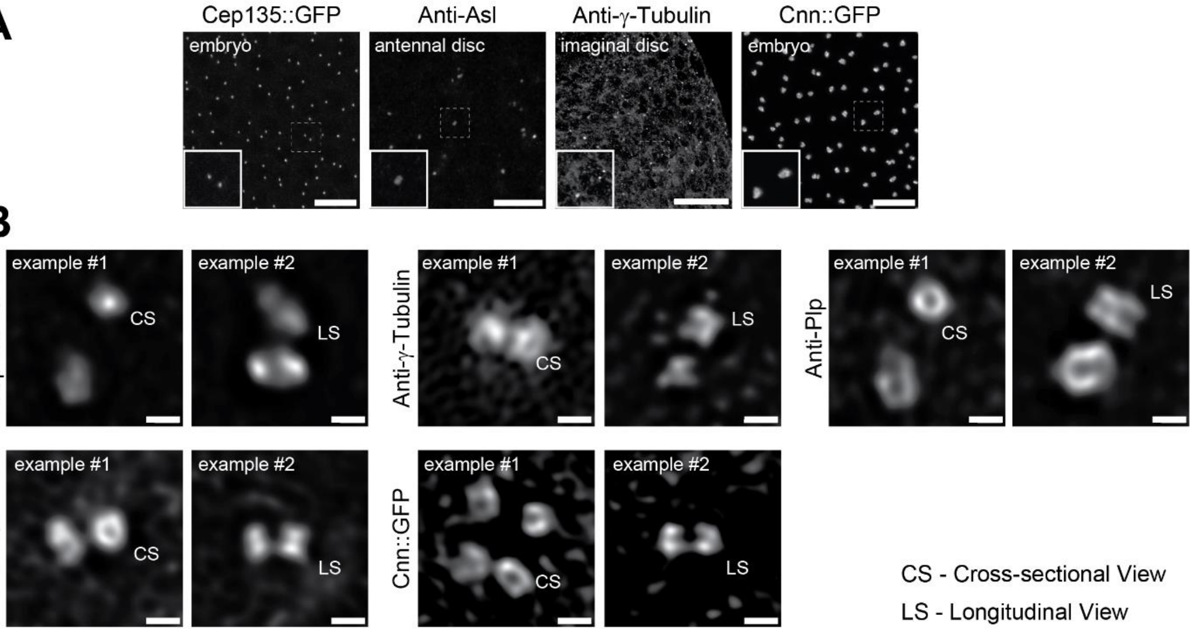

C *i*

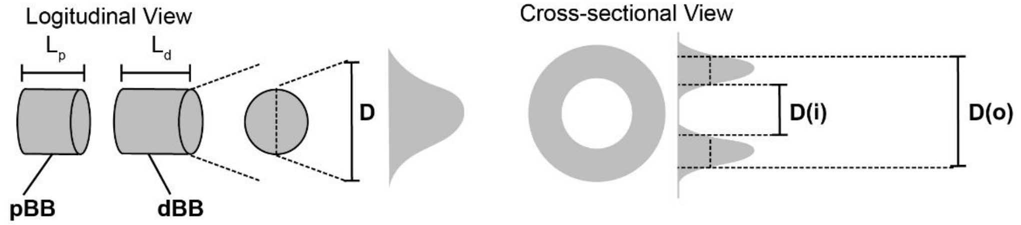

*ii*

| | proximal Basal Body (pBB) | | | | | distal Basal Body (dBB) | | | |
|---|---|---|---|---|---|---|---|---|---|
| | $L_p$ | $D_p$ | $D_p(i)$ | $D_p(o)$ | | $L_d$ | $D_d$ | $D_d(i)$ | $D_d(o)$ |
| **Cep135** | 178 ± 36 | 169 ± 17 | | | | 244 ± 38 | 283 ± 93 | | |
| **PACT** | U.D. | | 163 ± 32 | 360 ± 25 | | 234 ± 53 | | 215 ± 13 | 429 ± 26 |
| **γ-Tubulin** | U.D. | | 140 ± 35 | 281 ± 70 | | 284 ± 50 | | 190 ± 25 | 458 ± 39 |
| **CNN** | U.D. | | 164 ± 27 | 382 ± 30 | | 251 ± 34 | | 228 ± 08 | 456 ± 15 |
| **PLP** | U.D. | | 186 ± 26 | 380 ± 50 | | 363 ± 86 | | 279 ± 18 | 558 ± 36 |

*iii*

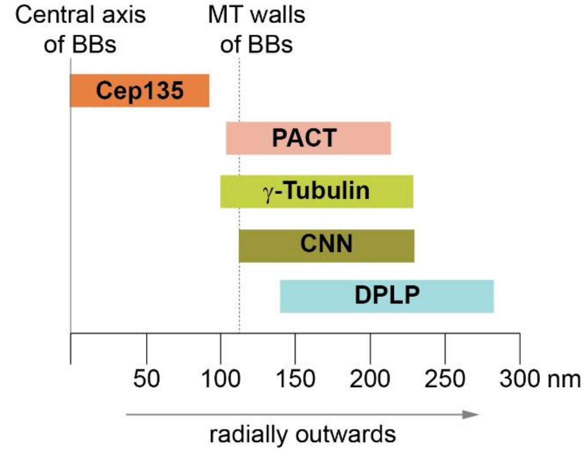

**Supplemental Figure 02: Localisation and quantification of the localisation patterns of centriole and PCM proteins in cycling cells and at the ciliary base in olfactory basiconic cilia in adult *Drosophila*.**

A) Representative confocal images of the centrosomal proteins (γ-tubulin, CNN, ASL, Cep135) in cycling cells of different *Drosophila* tissues, e.g., embryo, antennal disc, and imaginal disc. PACT is used as centriole wall marker. PCM proteins γ-tubulin and CNN are present in cycling cells (shown here) and at the ciliary base in OSNs (shown in Figure 2A(i)). On the other hand, ASL, another key PCM expansion factor, is present in cycling cells (shown here), but not detected at the ciliary base in OSNs (shown in Figure 2A(i)).

B) Representative Airyscan 2.0 images describe the localisation of four centrosomal proteins (γ-tubulin, CNN, PLP, and Cep135) and a centriole wall marker (PACT) in olfactory cilia.

C) Quantification of the localisation patterns of PCM components in OSNs obtained using Airyscan 2.0. Schemes show the method for quantification of proteins and different parameters (i). Length (mean±S.D.) and diameter (mean±S.D.) of the defined zones are mentioned in the table (ii). All values mentioned in the table are in nanometers (nm). n≥6 samples (U.D.=undetermined). S.D. - standard deviation. The schemes (in iii) represent the localisation patterns of the proteins drawn based on the quantification shown in (ii).

The experiments presented in A and B were repeated independently thrice. Scale bars on micrographs in A and B are 10 µm and 500 nm, respectively.

#### Supplemental Figure 03

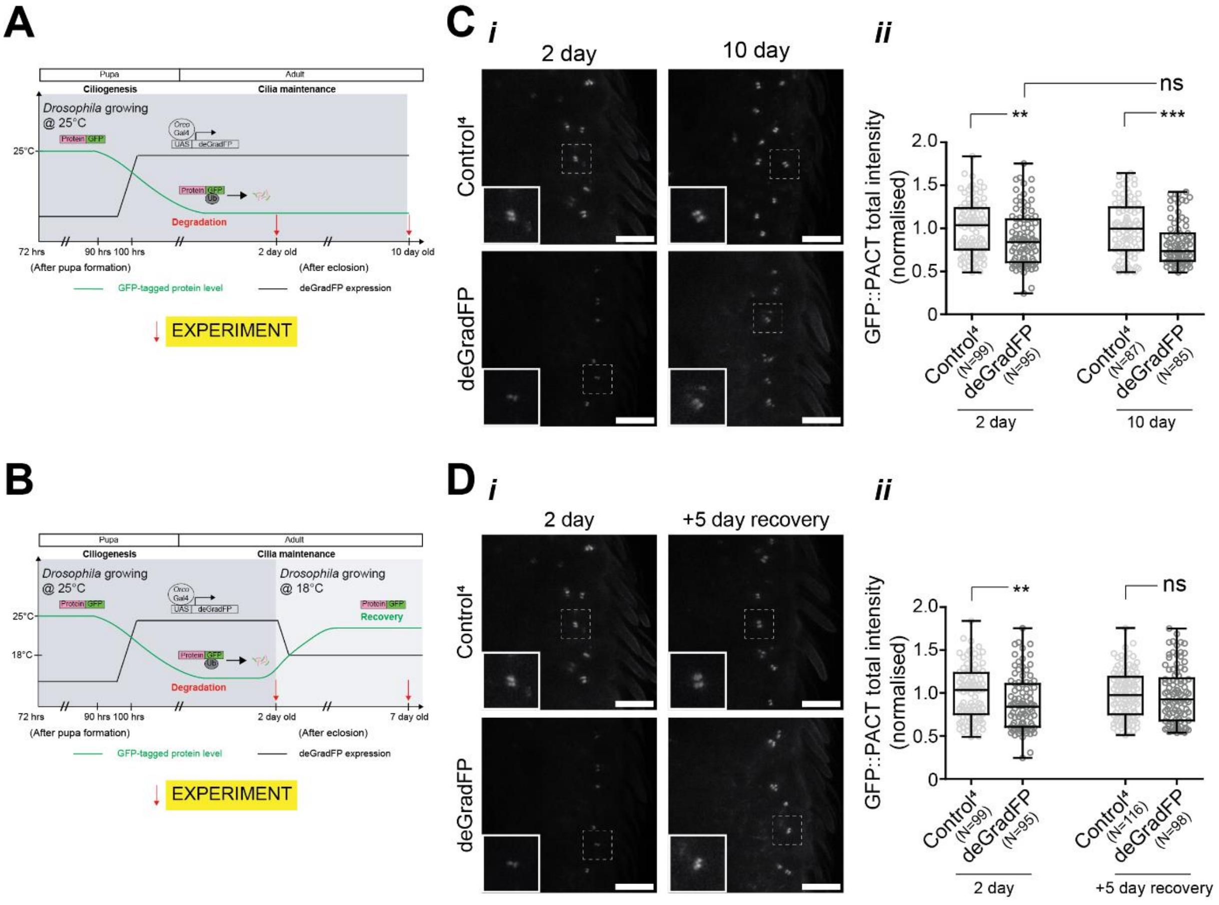

**Supplemental Figure 03: Centriole and PCM proteins are dynamically exchanged at the ciliary bases of basiconic OSNs.**

A, B) Scheme of the approach and timeline of the conditional degradation experiments. Scheme (A) represents the genetic strategy for temporally controlled degradation of GFP-tagged proteins in OSNs. Scheme (B) represents the genetic strategy for suppressing the degradation to study the recovery dynamics of GFP-tagged protein in OSNs.

C) Time-dependent changes in intensity of *pUbq*PACT::GFP (at 25°C) upon conditional degradation (under Gal4$^{Orco}$ driver) for 2 days and 10 days. Representative images (i) and their respective quantifications (ii). There is a significant reduction in *pUbq*PACT::GFP intensity upon expression of deGradFP for 2 days indicating that the deGradFP can target GFP-tagged protein for degradation in our system. Even after 10 days of deGradFP expression, the amount of GFP reduction does not increase significantly than that observed after 2 days of deGradFP expression. This suggests that the expression of new *pUbq*PACT::GFP protein occurs at a constant rate similar to the expression of *UAS*-deGradFP that is enough to compensate for the degradation of existing *pUbq*PACT::GFP protein.

D) Time-dependent changes in intensity of *pUbq*PACT::GFP (at 25°C) upon conditional degradation (under Gal4$^{Orco}$ driver) for 2 days and recovery (at 18°C) for 5 days post-degradation. Representative images (i) and their respective quantifications (ii).

Scale bars on confocal micrographs in C and D are 10 µm. All-range box plots of total intensity of GFP-tagged proteins at the base of olfactory basiconic cilia in the flies with specific genotypes and specific experimental condition are stated on the graphs.

### Supplemental Figure 04

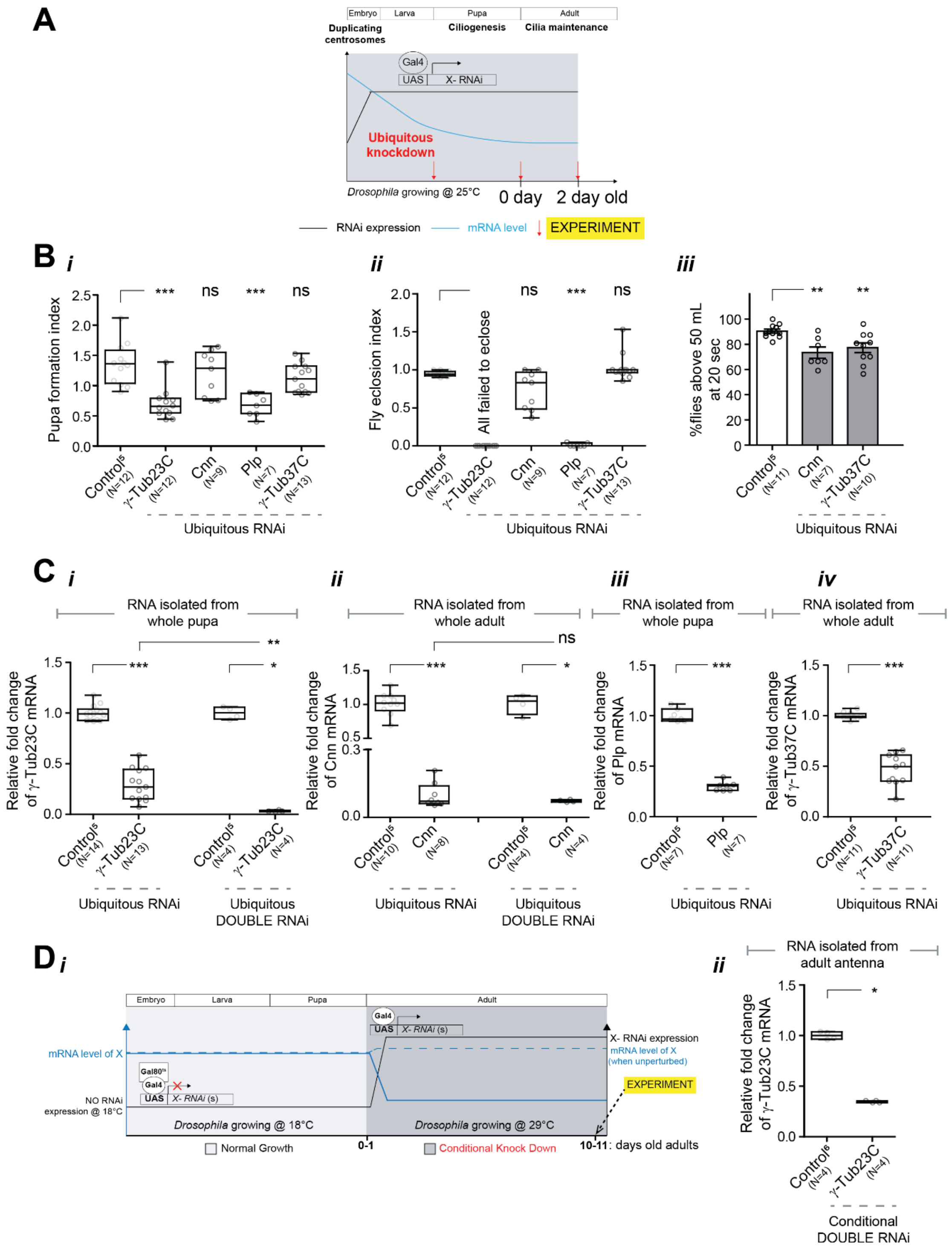

**Supplemental Figure 04: Validation of the efficacy of various dsRNA constructs used in PCM proteins knockdown.**

A) Scheme of the approach and timeline of the ubiquitous knockdown experiments.

B) Effect of ubiquitous knockdown of various PCM proteins in pupa formation (i) and pupa-to-adult fly eclosion (ii). Each cross gave two types of progeny due to presence of *UAS*-mCD8::GFP in the Gal4 driver line – GFP+ve progenies with knockdown and GFP-ve progenies with no knockdown (expected ratio of 1:1). γ-Tubulin23C and PLP knockdown showed ratio<1.0 indicating defects in larva-to-pupa conversion, however, there was no significant pupa formation defects in CNN knockdown. These observations phenocopy their respective null mutants[9–11]. In contrast, γ-Tubulin37C null mutants show embryonic lethality due to its role in female oocyte meiosis and early embryogenesis[12]. In our system, since Gal4$^{Tubulin}$ expression starts after embryogenesis, the RNAi-mediated depletion may occur too late to disrupt the early functions of γ-Tubulin37C leading to no defects in pupa formation and fly eclosion. We observe that all γ-Tubulin23C knockdown pupae fail to eclose. While very few of the PLP knockdown flies eclose, they are severely uncoordinated and die on the same day. Most of the CNN and γ-Tubulin37C knockdown pupae eclose. Negative geotaxis assay was performed on 2-day-old knockdown flies. Y-axis represents %flies to successfully climb half-height mark (50mL) at 20 sec after tapping (iii). CNN and γ-Tubulin37C knockdown flies show moderate negative geotaxis defects.

C) Relative change in γ-Tubulin23C (i), cnn (ii) Plp (iii) and γ-Tubulin37C expression (iv) in control and upon given candidate RNAi using Real-time qPCR keeping relevant housekeeping genes as normalisation factors. Note that all RNA values (Relative Quantification) were normalised using the formula (Relative Quantification = $2^{(-\Delta\Delta Ct)}$ and $\Delta\Delta Ct = \Delta Ct(Experiment) - \Delta Ct(Control)$).

D) Scheme of the approach and timeline of the conditional knockdown experiments (i). Relative change in γ-Tubulin23C expression in adult antennal tissue of control and γ-Tubulin23C DOUBLE RNAi using Real-time qPCR keeping relevant housekeeping genes as normalisation factors (ii).

### Supplemental Figure 05

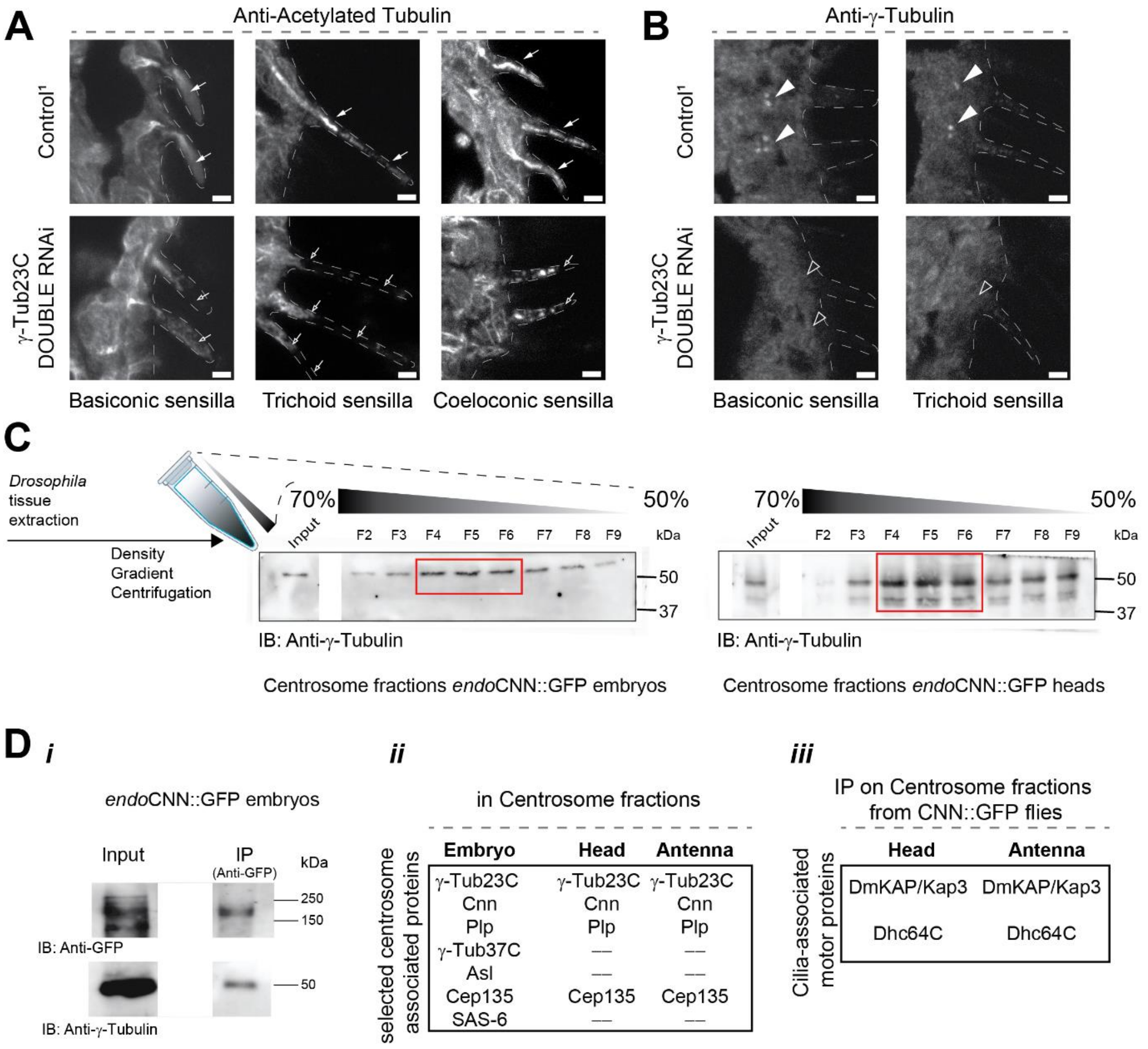

**Supplemental Figure 05: γ-Tubulin23C is essential for maintaining different types of mature olfactory cilia (A, B) and protein composition of centrosome/basal bodies purified from embryo and head/antenna is partly conserved (C, D).**

A) Acetylated-tubulin (ciliary marker) staining in basiconic, trichoid, and coeloconic olfactory cilia in control and γ-Tubulin23C conditional DOUBLE RNAi knockdown (under Gal4$^{Orco}$ driver) flies at 10 days in 29°C.

B) γ-tubulin staining at the ciliary base of olfactory cilia innervating basiconic and trichoid sensilla in control and γ-Tubulin23C conditional DOUBLE RNAi knockdown (under Gal4$^{Orco}$ driver) flies at 10 days in 29°C.

C) Scheme of the approach for protein identification in centrosomes purified from *Drosophila* cycling cells (embryo) and ciliated tissues (adult heads). γ-tubulin immunoblotting (IB) of centrosomal fractions purified from 2-to-4-hour-old embryos (left) and 0-to-2-day-old heads (right) with *endo*CNN::GFP. Centrosomal peaks for both tissues were detected in fractions 4-6 which were pooled for further analysis.

D) CNN (probed by anti-GFP) and γ-tubulin both are detected in embryo total protein lysate (Input, i). γ-tubulin is part of the co-IP complex of *endo*CNN::GFP isolated from 2-to-4-hour-old embryos (IP, i). List of centriole and PCM proteins found in isolated centrosomal fraction from 2-to-4-hour-old embryos, adult heads and antenna from 0-to-2-day-old flies (ii). Consistent with our protein localisation analysis (Figure 2A), ASL is not detected in *adult* Drosophila tissue. γ-Tubulin37C and SAS6 are also not detected as reported previously[4,12]. List of cilia-associated motor proteins found in co-IP complex of *endo*CNN::GFP from isolated centrosomal fractions (iii). Heterotrimeric kinesin-2 subunit KAP and cytoplasmic dynein heavy chain Dhc64C both were detected in adult heads and antennal tissue.

Scale bars on confocal micrographs in A and B are 2.5 µm each.

#### Supplemental Figure 06

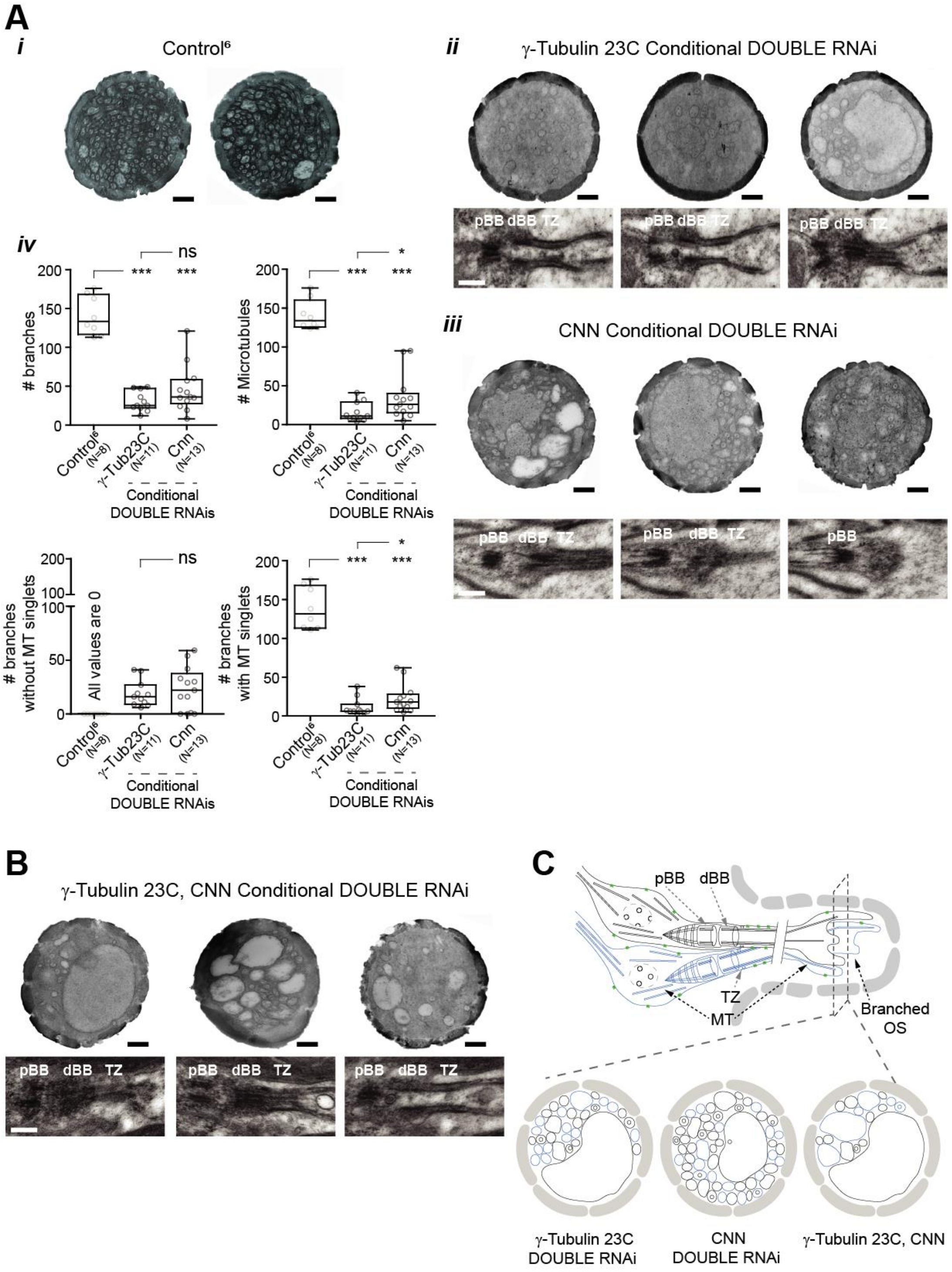

**Supplemental Figure 06: Levels of knock downs of γ-Tubulin23C and CNN are associated to defects observed on the ciliary shafts and γ-Tubulin23C and CNN synergistically work to maintain ciliary shaft.**

A) Representative electron micrographs of individual cross sections of branched OS from 10-day-old adult flies of control (i) and conditional DOUBLE RNAi knockdown of γ-Tubulin23C (ii, top) and CNN (iii, top), and respective branch quantifications (iv). Representative electron micrographs of individual longitudinal sections of BBs from 10-day-old adult flies of conditional DOUBLE knockdown of γ-Tubulin23C (ii, below) and CNN (iii, below).

B) Representative electron micrographs of individual cross sections of OS (top) and longitudinal (below) sections of BBs from 10-day-old flies with conditional simultaneous knockdown of γ-Tubulin23C with CNN, also shown and quantified in Figure 5E.

C) Scheme representing OSNs innervating a basiconic sensilla. Various features of the olfactory cilia - pBB and dBB, TZ, and OS branching are depicted for comparison with the micrographs. Scheme of OS cross-sections below demonstrates various observed differences in branching defects in conditional DOUBLE knockdowns of γ-Tubulin23C (left), CNN (middle) and conditional simultaneous knockdown of γ-Tubulin23C with CNN (right) highlighting their synergistic role in maintenance of ciliary shaft ultrastructure.

All electron micrographs in A and B represent features observed in n=3 samples (the experiments were repeated independently at least twice with similar results). Scale bars on all micrographs are 500 nm each.

### Supplemental Figure 07

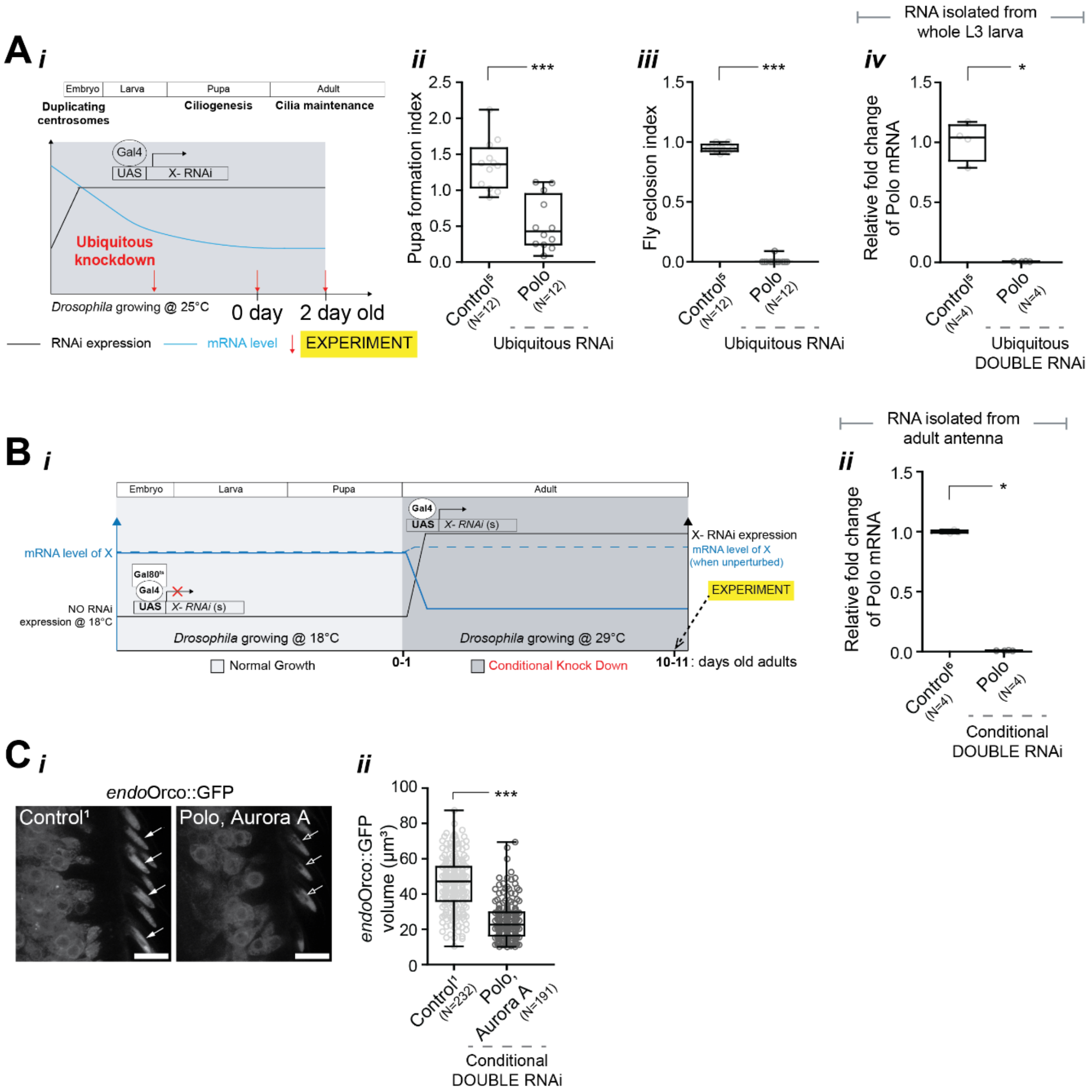

**Supplemental Figure 07: Validation of the efficacy of dsRNA constructs used in POLO knockdown**

A) Scheme of the approach and timeline of the ubiquitous knockdown experiments (i). POLO knockdown showed a pupa formation ratio <1.0 suggesting defects in larva-to-pupa conversion (ii). Furthermore, POLO knockdown pupae fail to eclose (iii). These observations phenocopy the null mutant as reported previously[13]. Relative change in POLO expression in control and upon POLO DOUBLE RNAi ubiquitous knockdown using Real-time qPCR keeping relevant housekeeping genes as normalisation factors (iv). Note that all RNA values (Relative Quantification) were normalised using the formula (Relative Quantification = $2^{(-\Delta\Delta Ct)}$ and $\Delta\Delta Ct = \Delta Ct(Experiment) - \Delta Ct(Control)$).

B) Scheme of the approach and timeline of the conditional knockdown experiments (i). Relative change in POLO expression in adult antennal tissue of control and upon POLO DOUBLE RNAi using Real-time qPCR keeping relevant housekeeping genes as normalisation factors (ii).

C) Time-dependent changes in volume of *endo*ORCO::GFP in the basiconic sensilla in control and upon conditional simultaneous knockdown of POLO with Aurora A for 10 days (at 29°C). Representative images (i) and respective quantifications (ii).

Scale bar on confocal micrographs in C are 10 µm.

#### Supplemental Figure 08

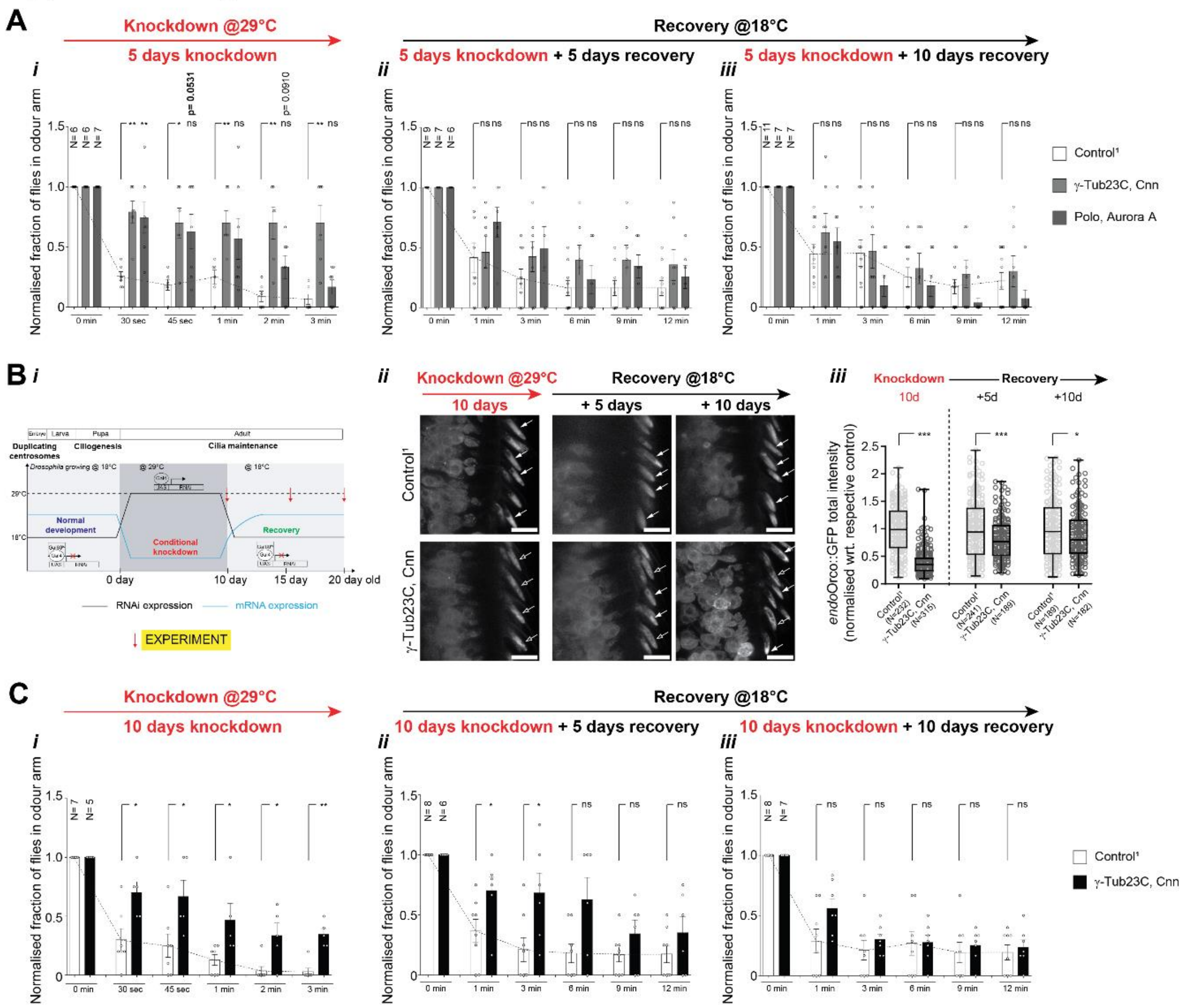

**Supplemental Figure 08: POLO and Aurora A work synergistically with PCM structural proteins at the ciliary base for maintenance of structure and function and regrowth of adult basiconic olfactory cilia.**

A) Time-dependent changes in the odour repulsion behaviours of control flies and flies with simultaneous conditional knockdown of γ-Tubulin23C with CNN and POLO with Aurora A for 5 days at 29°C (i) and recovery at 18°C for 5 days (ii) and 10 days (iii) post-knockdown. Each bar corresponds to a total of ≥60 flies measured in sets of 7-10 animals each. Note that we analysed behaviour of knockdown flies (kept in 29°C) until 3 min after odour exposure. However, during recovery (at 18°C), we observed that control flies showed sluggish response to repulsive odour possibly due to their slower physiological functioning at low temperature. Therefore, we analysed recovery behaviour over a longer timescale of 12 min after odour exposure.

B) Scheme of the approach and timeline of the conditional knockdown and recovery experiments (i). Time-dependent changes in *endo*ORCO::GFP intensity upon conditional knockdown for 10 days (at 29°C) and recovery (at 18°C). Representative images (ii) and respective quantifications (iii). ORCO intensities were normalised wrt. control of that condition for comparison since flies were grown at different temperatures for knockdown and recovery, which could affect the overall rate of protein synthesis and turnover.

C) Time-dependent changes in the odour repulsion behaviours of control flies and flies with simultaneous conditional knockdown of γ-Tubulin23C with CNN for 10 days at 29°C (i) and recovery at 18°C for 5 days (ii) and 10 days (iii) post-knockdown. Each bar corresponds to a total of ≥60 flies measured in sets of 7-10 animals each.

Scale bar on confocal micrographs in B are 10 µm.

#### References:

1. Jana, S. C., Mendonça, S., Werner, S. & Bettencourt-Dias, M. Methods to Study Centrosomes and Cilia in Drosophila. in *Cilia: Methods and Protocols* (eds Satir, P. & Christensen, S. T.) 215–236 (Springer, New York, NY, 2016). doi:10.1007/978-1-4939-3789-9_14.
2. Jana, S. C. *et al.* Kinesin-2 transports Orco into the olfactory cilium of Drosophila melanogaster at specific developmental stages. *PLOS Genet.* **17**, e1009752 (2021).
3. Chakraborty, T. S., Goswami, S. P. & Siddiqi, O. Sensory Correlates of Imaginal Conditioning in Drosophila melanogaster. *J. Neurogenet.* **23**, 210–219 (2009).
4. Jana, S. C. *et al.* Differential regulation of transition zone and centriole proteins contributes to ciliary base diversity. *Nat. Cell Biol.* **20**, 928–941 (2018).
5. Jana, S. C., Girotra, M., Ray, K. & Bettencourt, -Dias Monica. Heterotrimeric kinesin-II is necessary and sufficient to promote different stepwise assembly of morphologically distinct bipartite cilia in Drosophila antenna. *Mol. Biol. Cell* **22**, 769–781 (2011).
6. Werner, S. *et al.* IFT88 maintains sensory function by localising signalling proteins along Drosophila cilia. *Life Sci. Alliance* **7**, (2024).
7. Agarwal, R. G. *et al.* EB1 surges promote ciliary outer-segment growth through periodic tubulin influxes into the Drosophila olfactory cilia. *J. Cell Sci.* **139**, jcs263625 (2026).
8. Lehmann, V., Müller, H. & Lange, B. M. H. Immunoisolation of Centrosomes from Drosophila melanogaster. *Curr. Protoc. Cell Biol.* **29**, 3.17.1-3.17.13 (2005).
9. Sunkel, C. E., Gomes, R., Sampaio, P., Perdigão, J. & González, C. Gamma-tubulin is required for the structure and function of the microtubule organizing centre in Drosophila neuroblasts. *EMBO J.* **14**, 28–36 (1995).
10. Megraw, T. L., Kao, L.-R. & Kaufman, T. C. Zygotic development without functional mitotic centrosomes. *Curr. Biol.* **11**, 116–120 (2001).
11. Martinez-Campos, M., Basto, R., Baker, J., Kernan, M. & Raff, J. W. The Drosophila pericentrin-like protein is essential for cilia/flagella function, but appears to be dispensable for mitosis. *J. Cell Biol.* **165**, 673–683 (2004).
12. Tavosanis, G., Llamazares, S., Goulielmos, G. & Gonzalez, C. Essential role for γ-tubulin in the acentriolar female meiotic spindle of Drosophila. *EMBO J.* **16**, 1809–1819 (1997).
13. Sunkel, C. E. & Glover, D. M. Polo, a mitotic mutant of Drosophila displaying abnormal spindle poles. *J. Cell Sci.* **89**, 25–38 (1988).
